# An anaphase surveillance mechanism prevents micronuclei formation from mitotic errors

**DOI:** 10.1101/2021.02.26.433009

**Authors:** Bernardo Orr, Filipe De Sousa, Ana Margarida Gomes, Luísa T. Ferreira, Ana C. Figueiredo, Helder Maiato

## Abstract

Micronuclei are a hallmark of cancer and other human disorders and have recently been implicated in chromothripsis, a series of massive genomic rearrangements that may drive tumor evolution and progression. Here we show that Aurora B kinase mediates a surveillance mechanism that integrates error correction during anaphase with spatial control of nuclear envelope reformation to protect against micronuclei formation during human cell division. Using high-resolution live-cell imaging of human cancer and non-cancer cells we found that anaphase lagging chromosomes are often transient and rarely formed micronuclei. This strong bias against micronuclei formation relied on a midzone-based Aurora B phosphorylation gradient that assisted the mechanical transduction of spindle forces at the kinetochore-microtubule interface required for anaphase error correction, while delaying nuclear envelope reformation on lagging chromosomes, independently of microtubules. Our results uncover a new layer of protection against genomic instability and provide a strategy for the rational design of micronuclei-targeting therapies.

## Introduction

Micronuclei (MN) are small sized nuclei derived from chromosomes or chromosome fragments that fail to incorporate into daughter nuclei during cell division (Guo et al., 2019). MN formation is a widely used biomarker of cancer and inflammation, as well as several metabolic, reproductive, cardiovascular and neurodegenerative disorders (Fenech et al., 2020). Recently, MN derived from chromosome segregation errors have drawn exceptional attention due to their causal link with chromothripsis, a series of massive genomic rearrangements that may drive rapid tumor evolution and account for acquired drug resistance and oncogene activation (Crasta et al., 2012; Ly and Cleveland, 2017; Shoshani et al., 2020; Zhang et al., 2015). Chromosome segregation errors during mitosis are normally prevented before anaphase by a correction mechanism involving the kinase activity of the Chromosomal Passenger Protein (CPC) Aurora B at centromeres (Lampson and Grishchuk, 2017), under surveillance of the spindle assembly checkpoint (SAC) that monitors the establishment of kinetochore-microtubule (KT-MT) attachments (Musacchio, 2015). Despite this, between 0.1-10% of human primary/non-transformed or chromosomally stable cancer cells progress into anaphase with one or few chromosomes lagging behind due to merotelic attachments (Bakhoum et al., 2014; Cimini et al., 2002; Thompson and Compton, 2011; Worrall et al., 2018). Merotelic attachments, in which a single KT is attached to MTs from both spindle poles (Cimini et al., 2001), evade SAC detection and the resulting lagging chromosomes in anaphase constitute a potential source of MN (Crasta et al., 2012). However, whether anaphase lagging chromosomes inevitably result in MN remains controversial, with frequencies ranging from 18-78% reported in the literature (Cohen-Sharir et al., 2021; Fonseca et al., 2019; Huang et al., 2012; Thompson and Compton, 2011; Worrall et al., 2018). Importantly, despite that chromosomal instability (CIN) is a hallmark of human cancers (Bakhoum and Cantley, 2018; Hanahan and Weinberg, 2000), MN prevalence in both cancer and non-cancer cells is relatively infrequent (typically less than 5-10% of the cells) (Bonassi et al., 2011; Jdey et al., 2017), even after induction of massive chromosome segregation errors by experimental abrogation of the SAC (Cohen-Sharir et al., 2021), implying the existence of surveillance mechanisms that either prevent MN formation from SAC-invisible mitotic errors or account for the clearance of micronucleated cells. While mechanisms of MN prevention from mitotic errors remain unknown, clearance of micronucleated cells involves a p53-dependent mechanism that causes cell cycle arrest/apoptosis (Fonseca et al., 2019; Janssen et al., 2011; Sablina et al., 1998; Thompson and Compton, 2010) and an innate immune response mediated by cGAS-STING, which is thought to sense cytosolic DNA on MN with ruptured nuclear envelopes (NEs) (Mackenzie et al., 2017; Santaguida et al., 2017). Given their potential role in the genesis of MN and the respective implications for human health, here we investigated how human cells deal with SAC-invisible mitotic errors that result in anaphase lagging chromosomes. Our findings uncover an active surveillance mechanism operating during anaphase that protects against MN formation from mitotic errors.

## Results

### Anaphase lagging chromosomes rarely form micronuclei

Many intrinsic factors may influence whether a lagging chromosome results in a MN. This appears to vary among different chromosomes (Worrall et al., 2018) and depend on their delay/position relative to the main segregating chromosome mass (Fonseca et al., 2019) or the presence/absence of centromeres (Norppa and Falck, 2003). In addition, external experimental factors, such as the spatiotemporal resolution of the microscopy setup/analysis and sample size, may undermine our capacity to accurately determine the fate of lagging chromosomes in living cells. To mitigate these external limitations, we used 4D live-cell spinning-disk confocal microscopy covering the entire chromosome set and the mitotic spindle, with 30 sec temporal resolution, to determine the fate of anaphase lagging chromosomes in human cells. This allowed the unequivocal identification of all lagging chromosomes, including those of highly transient nature that normally resolve early in anaphase. We found that 9% of chromosomally stable (non-transformed) RPE1 cells and 44% of chromosomally unstable (transformed) U2OS cells displayed at least one transient lagging chromosome during anaphase (Fig. 1a, b; Fig. S1a; Movies S1-2). However, only 6% and 14% of the lagging chromosomes in RPE1 and U2OS cells, respectively, resulted in MN (Fig. 1a, Fig. S1a, c; Movies S1-2). Thus, lagging chromosomes in both transformed and non-transformed human cells are often corrected during anaphase, and show a strong bias to re-integrate the main nuclei.

**Fig. 1.**
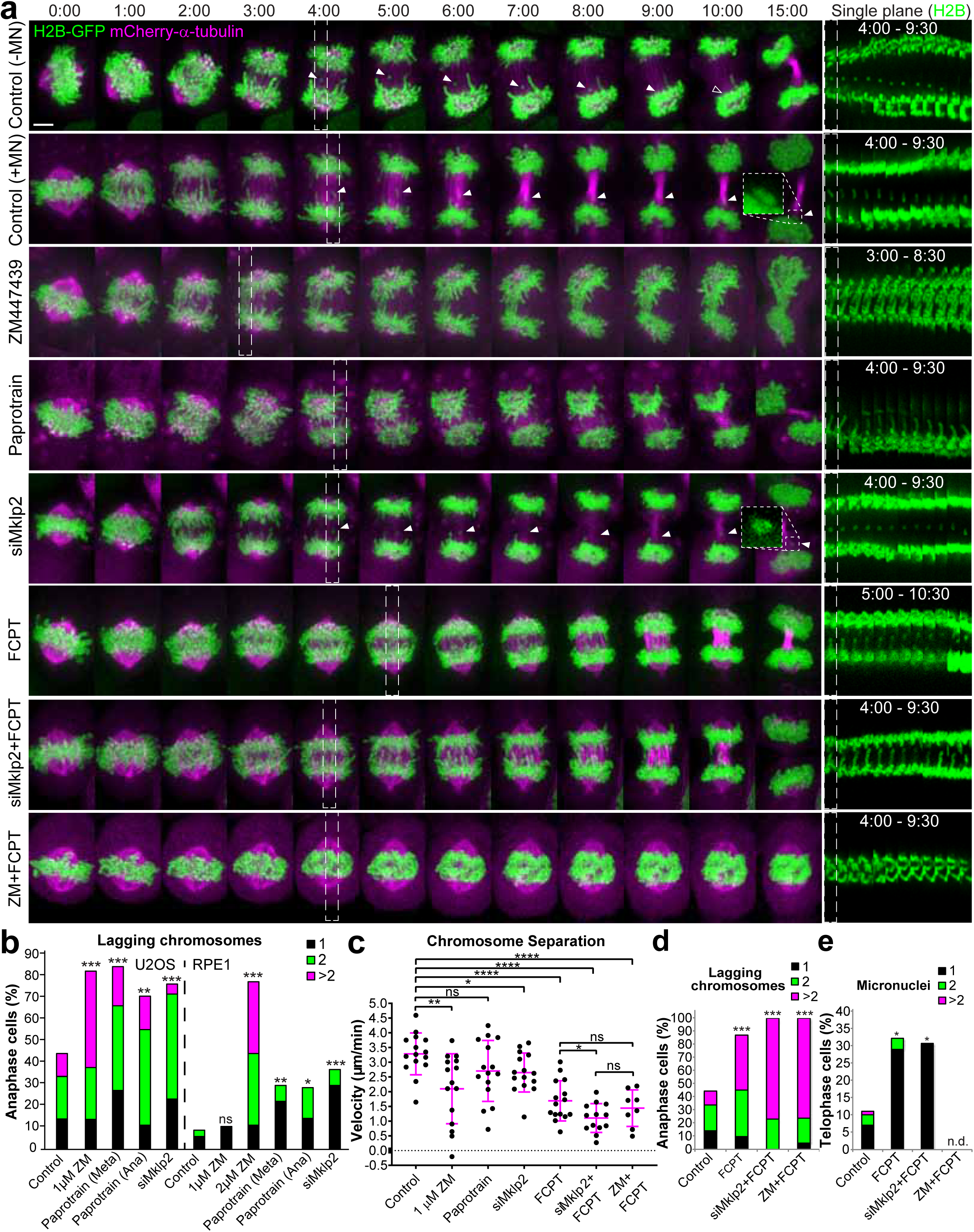
Anaphase lagging chromosomes are frequent, but normally corrected by a midzone-based Aurora B activity gradient that assists spindle forces to prevent micronuclei formation. **a**, U2OS cells stably expressing H2B-GFP (H2B) and mCherry-α-tubulin under specified conditions. White dashed rectangular ROIs indicate the first frame shown in the single plane kymographs on the right for each condition. White arrowheads track lagging chromosomes until their eventual re-integration in the main nuclei or form MN. Higher magnification (3X) insets show single channel (H2B) examples of MN. Scale bar = 5 μm. **b**, Frequency of U2OS and RPE1 cells with anaphase lagging chromosomes under the specified conditions. U2OS cells: [Control n=198, 1 µM ZM447439 (ZM) n=29, 10 µM Paprotrain added in Metaphase (Meta) n=33, 10 µM Paprotrain added in Anaphase (Ana) n=45, siMklp2 n=43]. RPE1 cells: [Control n=136, 1 µM ZM447439 (ZM) n=19, 2 µM ZM447439 (ZM) n=18, 10 µM Paprotrain added in Metaphase (Meta) n=27, 10 µM Paprotrain added in Anaphase (Ana) n=21, siMklp2 n=27]. **c,** Chromosome separation velocity for U2OS cells under specified conditions (magenta lines indicate mean±SD). Control n=15, 1 µM ZM (ZM447439) n=15, Paprotrain (10 µM added in Anaphase) n=15, siMklp2 n=15, FCPT (100 µM) n=15, siMklp2+FCPT (100 µM) n=13; ZM(1 µM)+FCPT(100 µM) n=8. Frequency of U2OS cells with **d**, lagging chromosomes and **e**, MN for the specified conditions (Control n=198, 100 µM FCPT n=31, siMklp2 + 100 µM FCPT n=13, 100 µM FCPT + 1 µM ZM n=21).

### Anaphase errors are frequent but are normally corrected by a mechanism relying on a midzone-based Aurora B phosphorylation gradient

In anaphase, Aurora B transfers from centromeres to the spindle midzone where it forms a phosphorylation gradient (Afonso et al., 2014; Fuller et al., 2008). To investigate whether Aurora B plays a role in anaphase error correction, we performed acute inhibition of Aurora B activity with ZM447439 after anaphase onset (Fig. 1a, b; Fig. S1a; Fig. S2a; Movies S1-3). In parallel, we inhibited Aurora B transport to the spindle midzone either by acute inhibition of Mklp2/kinesin-6 after anaphase onset with Paprotrain (Tcherniuk et al., 2010) or constitutive Mklp2 depletion by RNAi (Gruneberg et al., 2004) (Fig. 1a, b; Fig. 2a, b; Fig. S1a; Fig. S2b-e, Fig. S3a-d; Movies S1-3). While ZM447439 treatment did not affect the re-localization dynamics of Aurora B to the spindle midzone, this was significantly delayed or abolished after Paprotrain treatment or Mklp2 knockdown, respectively (Fig. 2a, b; Movie S3). All conditions abolished the formation of a phosphorylation gradient on both segregating and lagging chromosomes (Fig. 3a-c), while significantly increasing the frequency of anaphase cells with lagging chromosomes and the number of lagging chromosomes per cell in both RPE1 and U2OS cells (Fig. 1a, b; Fig. S1a; Movies S1-2). Noteworthy, resolution of anaphase DNA bridges depended on Aurora B activity but not on its midzone localization (Fig. S1b). Thus, most anaphase cells normally have one or more transient lagging chromosomes that are actively corrected by a mechanism relying on the establishment of a midzone-based Aurora B phosphorylation gradient.

**Fig. 2.**
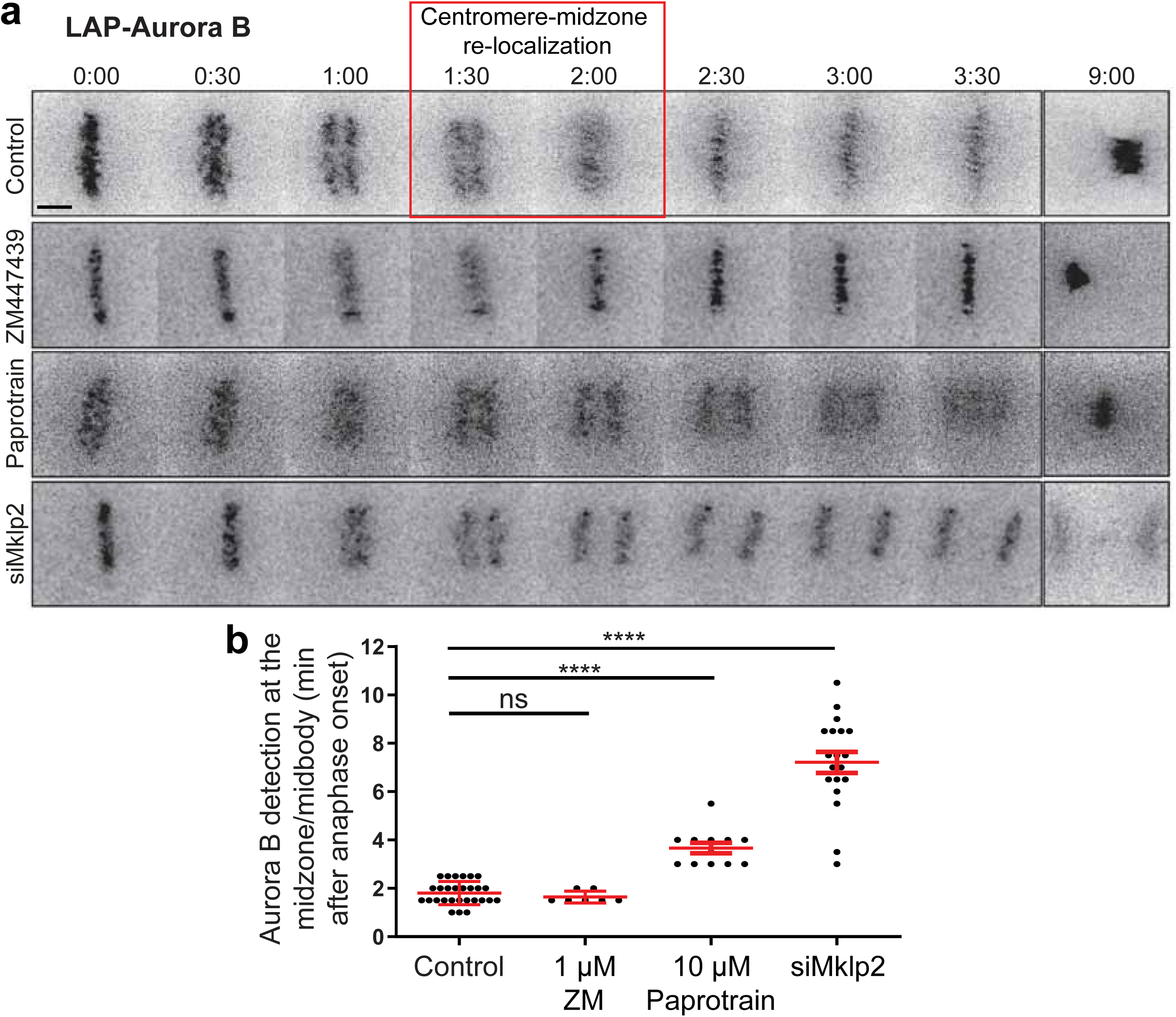
Mklp2 is required for dynamic Aurora B re-localization from centromeres to the spindle midzone. **a**, Selected time-frames of representative HeLa cells stably expressing LAP-Aurora B in control, 1 μM ZM447439 (ZM) treated, 10 µM Paprotrain (Papro) treated or after depletion of Mklp2. Time = min:sec. ZM and Paprotrain treatments were added within the first min of anaphase onset. Red box highlights the narrow time window of centromere to midzone re-localization in control cells. Scale bar = 5 µm. **b**, Quantification of the time it takes to detect Aurora B at the midzone/midbody after anaphase onset (Control n=28, ZM n=7, Paprotrain n=14, siMklp2 n=19).

**Fig. 3.**
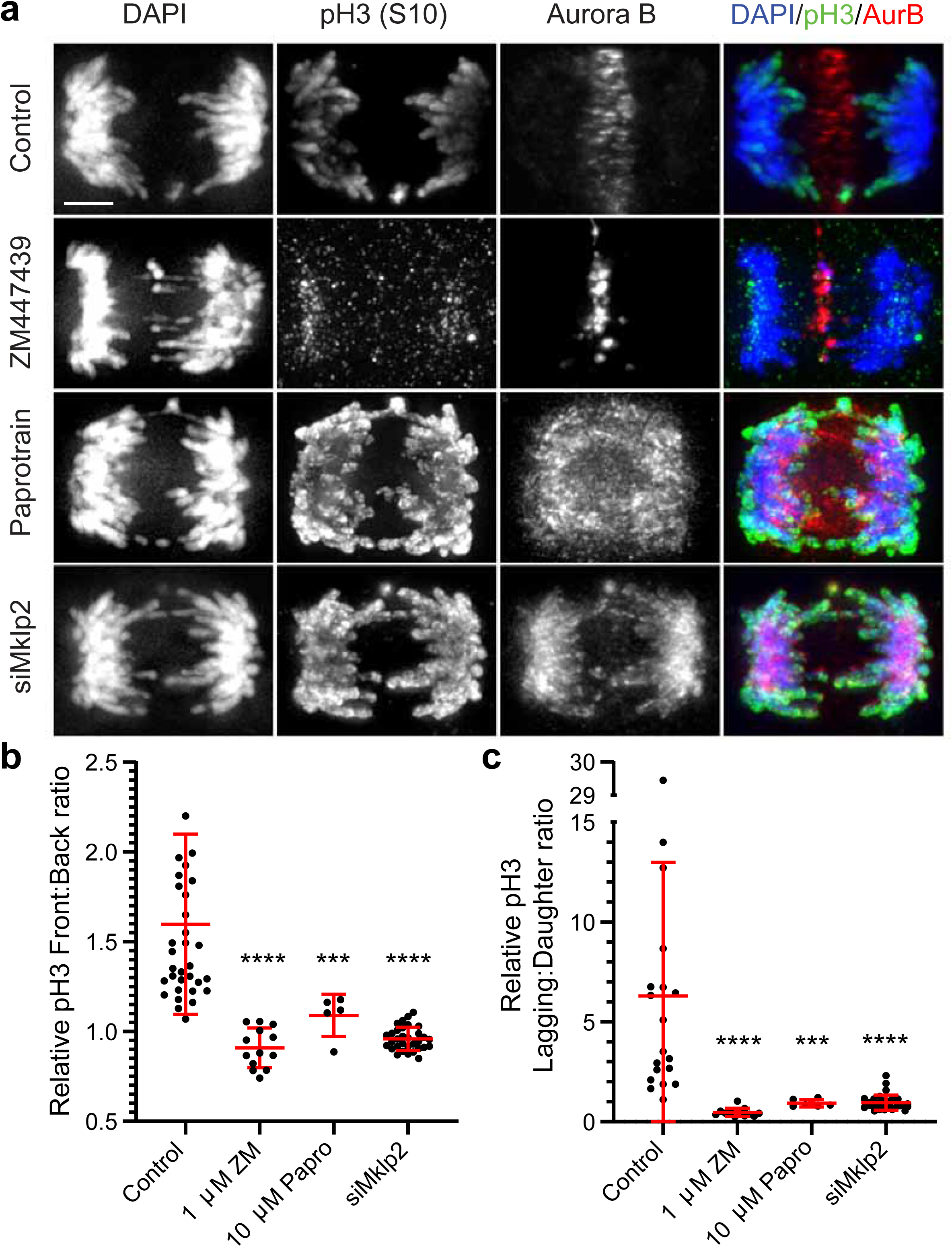
An Aurora B activity gradient on chromosomes during anaphase is dependent on its midzone localization in human cells. **a**, Representative images of U2OS cells fixed and stained to reveal DAPI (blue), pH3 Ser10 (green) and Aurora B (red). siMklp2 treated cells were imaged 72 h after siRNA transfection and treatments with ZM447439 and Paprotrain were performed for 7 min before fixing and processing cells for immunofluorescence. Scale bar = 5 µm. **b**, Relative Front/Back pH3 Ser10 ratio of U2OS anaphase cells calculated for control cells (n=34), 1 µM ZM447439 (n=13), 10 µM Paprotrain (n=5) and siMklp2 (n=32). **c**, Relative Lagging/Daughter pH3 Ser10 ratio of U2OS anaphase cells calculated for control cells (n=19), 1 µM ZM447439 (n=13), 10 µM Paprotrain (n=6) and siMklp2 (n=33).

### Midzone Aurora B-mediated error correction during anaphase protects against MN formation

Next, we determined whether Aurora B-mediated error correction during anaphase protects against MN formation. Global inhibition of Aurora B activity in anaphase impacted chromosome separation by affecting spindle midzone organization and elongation, as well as chromosome segregation distance (Fig. 1a, c; Fig. S4a, b). There was also a massive increase in the frequency of DNA bridges, which, upon decondensation, often resulted in the formation of a continuous chromatin mass with the other partially segregated chromosomes (Fig. 1a; Fig. S1b; Movies S1-2). Paprotrain treatment after anaphase onset delayed, but did not fully prevent Aurora B localization at the spindle midzone, yet it strongly reduced spindle elongation velocity (Fig. 2a, b; Fig. S4a). Importantly, the frequency of cells with lagging chromosomes that resulted from acute Paprotrain treatment in metaphase or anaphase were indistinguishable from the one observed after constitutive Mklp2 depletion (Fig. 1b), suggesting an anaphase-specific role for a midzone-based Aurora B activity gradient in error correction. Therefore, we concentrated on investigating MN formation from mitotic errors after Mklp2 depletion. The frequency of telophase cells with MN increased two- and five-fold in U2OS and RPE1 cells, respectively (Fig. S1c). Moreover, the probability of lagging chromosomes resulting in MN also increased in both cell types (Fig. S1c). Thus, the observed increase in MN did not simply reflect the increase in lagging chromosomes after Mklp2 depletion, suggesting that a midzone-based Aurora B activity gradient is required to prevent MN formation beyond its role in anaphase error correction.

### Midzone Aurora B mediates anaphase error correction by assisting the mechanical transduction of spindle forces at the kinetochore-microtubule interface

Lagging chromosomes may be corrected during anaphase by the mechanical action of spindle forces that ultimately resolve merotelic attachments (Cimini et al., 2004; Cimini et al., 2003). However, the underlying molecular mechanism remains unclear. In yeast, spindle elongation relies on the functional interaction between the CPC and kinesin-5 (Rozelle et al., 2011). To investigate whether Aurora B mediates anaphase error correction by regulating spindle forces acting at the KT-MT interface, we attenuated spindle elongation in U2OS cells with FCPT, which prevents interpolar MT sliding by locking kinesin-5 on MTs (Collins et al., 2014), and compared its effects with those observed after the disruption of Aurora B function in anaphase. Acute FCPT treatment at anaphase onset strongly impaired spindle elongation, chromosome separation velocity and chromosome segregation distance (Fig. 1a, c; Fig. S4a, b; Movie S1). A similar effect was observed after acute Aurora B inhibition at anaphase onset or Mklp2 depletion by RNAi, albeit to a lesser extent in the latter (Fig. 1a, c; Fig. S4a, b; Movie S1). Combination of FCPT treatment with acute Aurora B inhibition at anaphase onset or Mklp2 depletion only marginally impaired spindle elongation and chromosome separation beyond the effect caused by FCPT treatment alone (Fig. 1a, c; Fig. S4a, b; Movie S1). In agreement, we observed an equivalent increase in the frequency of anaphase cells with lagging chromosomes and a consequent increase in telophase cells with MN, either with FCPT alone or combined with acute Aurora B inhibition or Mklp2 depletion, in line with our previous observations after Aurora B inhibition or Mklp2 depletion alone (Fig. 1a, d, e; Movie S1). These data suggest that a midzone-based Aurora B activity gradient mediates anaphase error correction by assisting the transduction of spindle forces at the KT-MT interface.

### Aurora B substrates involved in stabilization of kinetochore-microtubule attachments are required for anaphase error correction and micronuclei prevention

To investigate whether anaphase error correction requires the establishment of stable KT-MT attachments we performed a high-throughput mini-screen in live HeLa cells after RNAi-mediated depletion of KMN network components, including several Aurora B substrates (Cheeseman et al., 2006; Welburn et al., 2010), and Aurora B itself. Focusing exclusively on chromosomes that lagged behind in anaphase after completing alignment to the metaphase plate, we found that their correction and subsequent re-integration into the main nuclei were greatly compromised by any experimental condition that disrupted KT-MT attachment stability (Fig. 4a-c). Moreover, the outcome of Aurora B depletion by RNAi was more benign than by acute inhibition (likely due to residual kinase activity in the former) and was indistinguishable from the depletion of KMN network components (Fig. 4a-c). These data suggest that, contrary to promoting MT detachment from KTs during error correction in early mitosis when it localizes at centromeres (Lampson et al., 2004), Aurora B stabilizes KT-MT attachments once it transfers to the spindle midzone in anaphase.

**Fig. 4.**
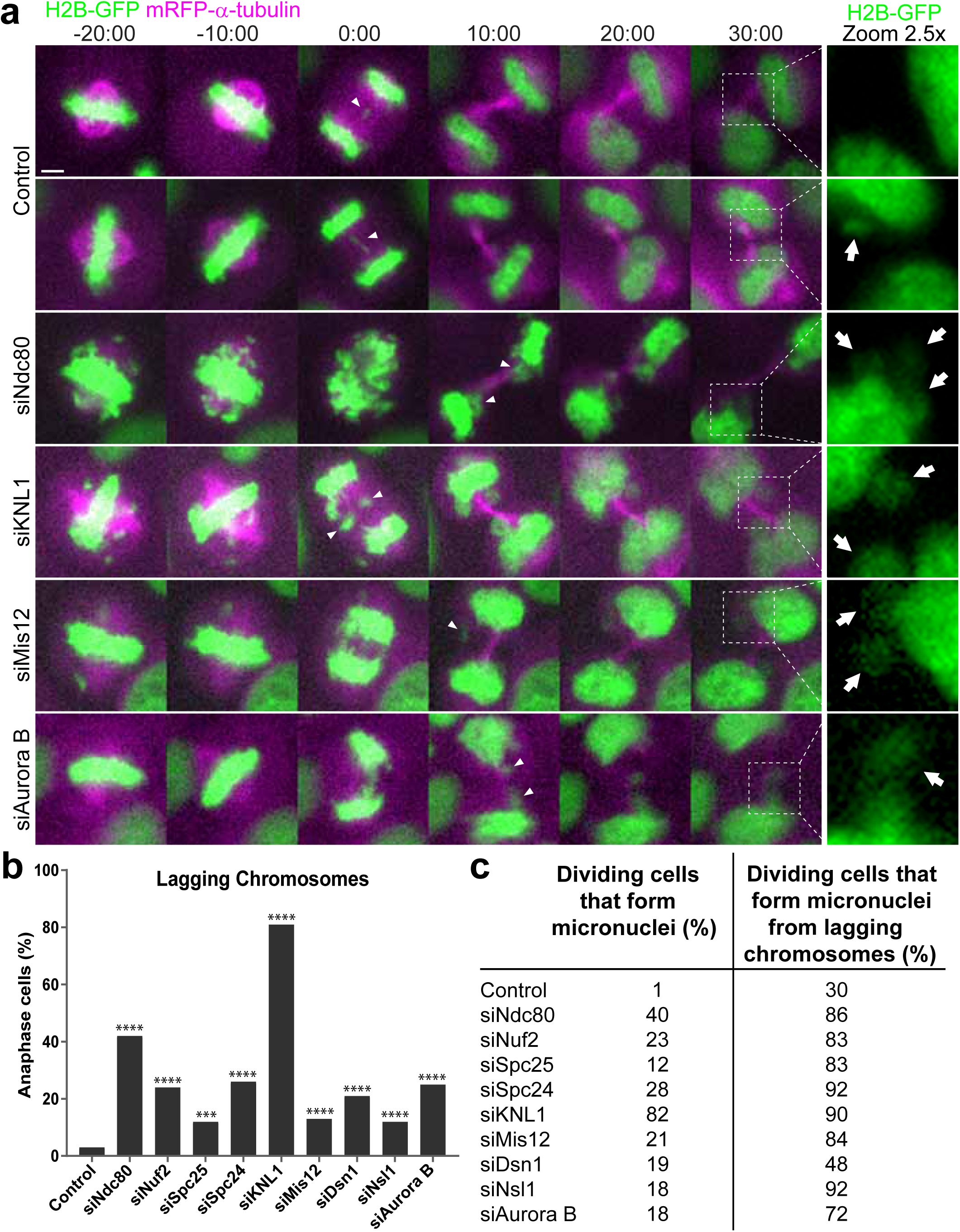
Anaphase error correction and micronuclei prevention requires stable kinetochore-microtubule interactions. **a**, HeLa cells stably expressing H2B-GFP and mRFP-α-tubulin treated with scramble siRNA (control) or siNdc80, siKNL1, siMis12. Time=min:sec. Time 00:00 =anaphase onset. White arrowheads indicate lagging chromosomes and arrows show MN. Scale bar=5 μm. **b,** Frequency of anaphase cells with lagging chromosomes for all conditions. **c,** Frequency of dividing cells that form MN (left column) and of dividing cells with lagging chromosomes that form MN (right column). Control n=355, siNdc80 n=102, siNuf2 n=100, siSpc25 n=99, siSpc24 n=100, siKNL1 n=194, siMis12 n=200, siDsn1 n=459, siNsl1 n=200, siAurora B n=287 cells.

### A midzone Aurora B phosphorylation gradient, rather than midzone microtubules, spatially controls NER on anaphase lagging chromosomes

We have previously proposed a ‘chromosome separation checkpoint’ in which a midzone-based Aurora B activity gradient monitors the extent of chromosome separation during anaphase to spatially control nuclear envelope reformation (NER) (Afonso et al., 2014; Maiato et al., 2015). Based on these findings, one may predict that a delay of NER on lagging chromosomes would allow anaphase error correction and open a window of opportunity for their subsequent re-integration in the main segregating chromosome mass, in which the recruitment of non-core NE components is also delayed in the midzone-facing side (Gerlich and Ellenberg, 2003; Lu et al., 2011). To test this hypothesis, we started by monitoring the Aurora B phosphorylation gradient on chromosomes using a rabbit monoclonal antibody against phospho-histone H3 (pH3) on Ser10 in fixed cells, after induction of chromosome segregation errors in non-transformed RPE1 cells by performing a 6 h nocodazole treatment/washout (Liu et al., 2018). However, we found that this procedure significantly perturbed the Aurora B phosphorylation gradient in anaphase (Fig. S5a-c), to an extent equivalent to Mklp2 depletion in human cells (Fig. 3a-c). To overcome this limitation, we implemented a high-resolution live-cell microscopy assay to simultaneously image the entire chromosome set labelled either with H2B-mRFP, or a Cy3-conjugated Fab fragment against pH3 on Ser10 (Hayashi-Takanaka et al., 2009), and two GFP-tagged non-core NE proteins (Nup153 and LBR) in human U2OS cells, which show an intrinsic high rate of chromosome segregation errors in anaphase. We found that, in striking contrast with controls, the majority of pH3-positive lagging chromosomes and DNA bridges in Mklp2-depleted cells recruited Nup153 and LBR (Fig. 5a-d, g; Fig. 6a-d; Fig. S6a, b; Fig. S7a-c; Movies S4-6). Quantitative analyses revealed that as lagging chromosomes in control cells were gradually corrected and moved away from the spindle midzone during anaphase, they began recruiting Nup153 and LBR after a significant delay relative to the main segregating chromosome masses, and this delay was dependent on the establishment of a midzone-based Aurora B phosphorylation gradient (Fig. 6e; Fig. S7d; Movies S5-6). Thus, NE defects on lagging chromosomes are often transient and thus may not inevitably result in pathological conditions. Interestingly, Mklp2-depleted cells showed normal Nup153 and LBR recruitment to the main daughter nuclei, despite retaining pH3 for extended periods (Fig. 5e, f; Fig. 6f; Fig. S7e; Movies S5-6). This indicates that pH3 is not the signal that determines the recruitment of non-core NE proteins, in line with our previous findings in *Drosophila* cells (Afonso et al., 2014). However, a clear correlation between Aurora B activity on lagging chromosomes and Nup153 recruitment was observed in control cells (Fig. 5h) and this was dependent on Aurora B localization at the spindle midzone (Fig. 5i). Lastly, to exclude the possible role of MTs in selectively preventing the recruitment of non-core NE components (Liu et al., 2018; Liu and Pellman, 2020) we simultaneously monitored Aurora B activity on chromosomes, Nup153 recruitment, and the distribution of spindle MTs in the same cell over time, with or without Mklp2. We found that Nup153 recruitment to lagging chromosomes after Mklp2 RNAi could not be explained by the disassembly of midzone MTs, as these were clearly present and found to co-localize with MN that recruited Nup153 (Fig. 7a, b; Movie S7). Quantitative super-resolution CH-STED analysis in fixed cells (Pereira et al., 2019) confirmed our live-cell data, since we could not detect differences in the density of MT bundles surrounding anaphase lagging chromosomes, with or without Mklp2 (Fig. 7c, d). We concluded that a midzone-dependent Aurora B activity gradient on chromosomes, rather than midzone MTs, is required to delay the completion of NER on lagging chromosomes in human cells to prevent MN formation.

**Fig. 5.**
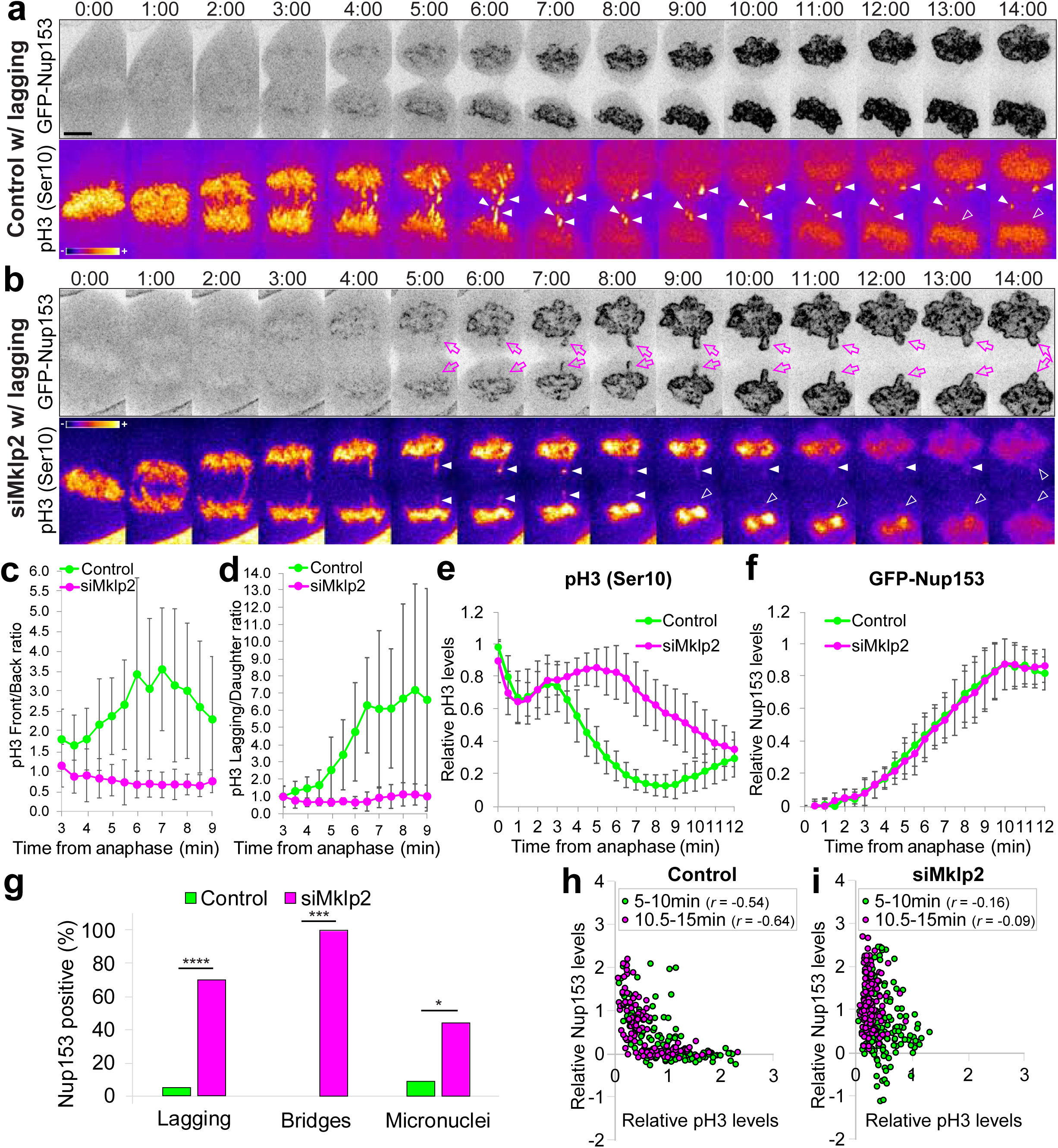
A midzone-based Aurora B activity gradient spatially regulates the completion of nuclear envelope reassembly on anaphase lagging chromosomes in human cells. **a**, Control cell with lagging chromosomes that are negative for Nup153 (inverted, grayscale) and positive for pH3 (Fire LUT). **b,** Mklp2-depleted cell with lagging chromosomes that are positive for both Nup153 (inverted, grayscale) and pH3 (Fire LUT). Time=min:sec. White arrowheads track lagging chromosomes until their eventual re-integration in the main nuclei or form MN. Magenta arrows highlight lagging chromosomes that prematurely accumulate Nup153 in siMklp2-depleted cells. Scale bar = 5 μm. Quantification of pH3 Ser10 **c,** Front/Back and **d,** Lagging/Daughter ratio on segregating chromosomes during anaphase. Control n=10; siMklp2 n=10; error bars represent SD. Quantification of **e,** pH3 Ser10 and **f,** GFP-Nup153 on daughter chromosomes/nuclei in control and siMklp2-treated cells. Control n=20; siMklp2 n=20; error bars represent SD. **g,** Frequency of mitotic errors that are positive for Nup153 in control and siMklp2-depleted cells (Control n=126 lagging, n=5 bridges, n=23 MN; siMklp2 n=120 lagging, n=4 bridges, n=18 MN). Correlation between the relative levels of Nup153 and pH3 on lagging chromosomes for **h,** Control and **i,** Mklp2-depleted cells. The calculated Pearson correlation coefficient (*r*) is indicated for each condition and time interval.

**Fig. 6.**
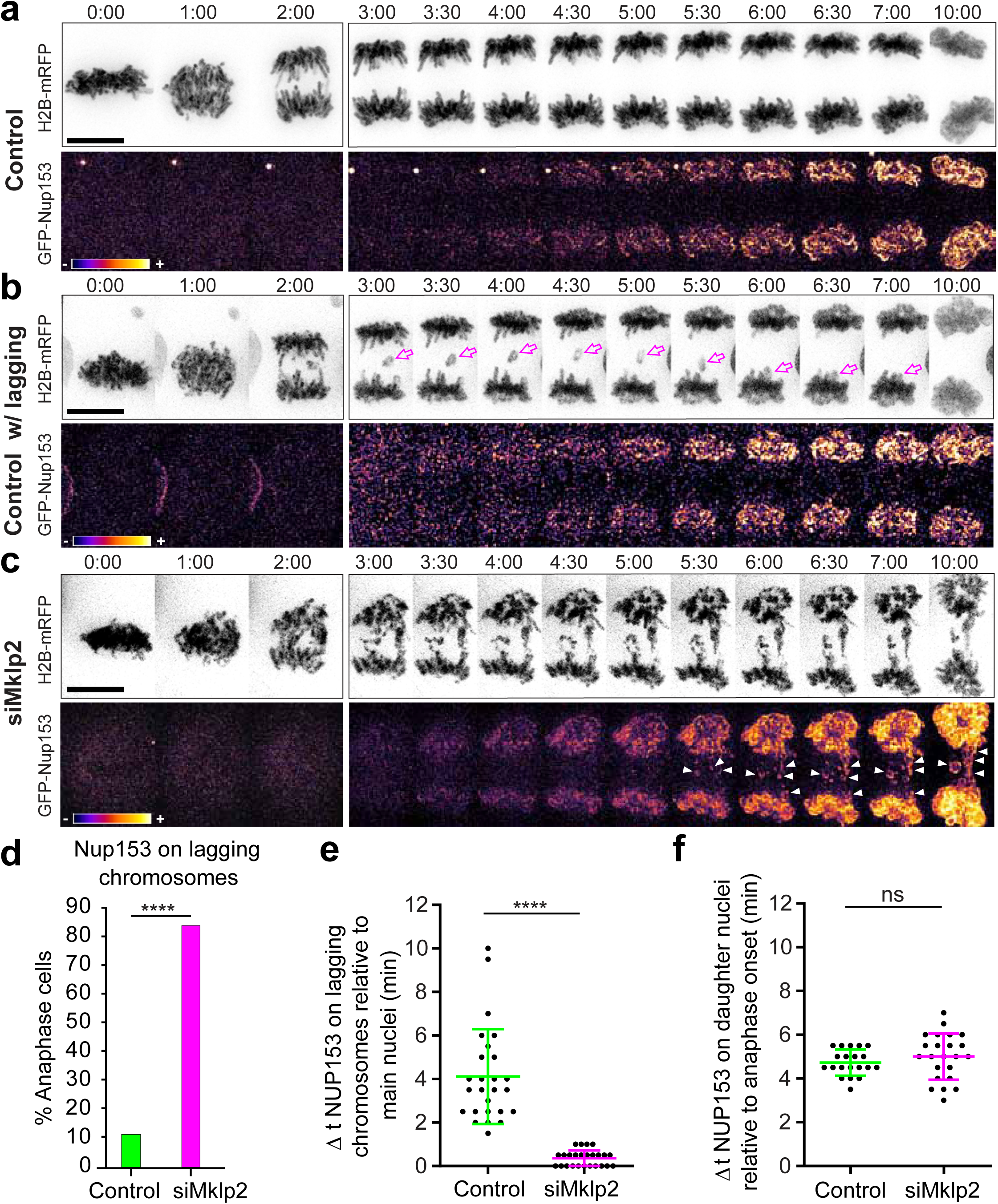
Aurora B midzone localization prevents premature Nup153 accumulation on lagging chromosomes. Representative examples of **a,** control cell without errors, **b,** control cell with a lagging chromosome and **c,** Mklp2-depleted cell with segregation errors. Time is shown in min:sec. Cells shown in panels (A-C) are U2OS cells stably expressing mRFP-H2B (inverted; grayscale) and transiently expressing eGFP-Nup153 (FIRE LUT). Scale bar = 5 μm. **d,** Quantification of the percentage of anaphase cells (Control n=26, siMklp2 n=25) in which Nup153 was detected on lagging chromosomes. **e,** Difference in time (min) between detection of Nup153 on daughter nuclei and lagging chromosomes (Control n=26, siMklp2 n=25). **f,** Quantification of the time (min) it takes to detect Nup153 on daughter nuclei after anaphase onset (Control n=45, siMklp2 n=30).

**Fig. 7.**
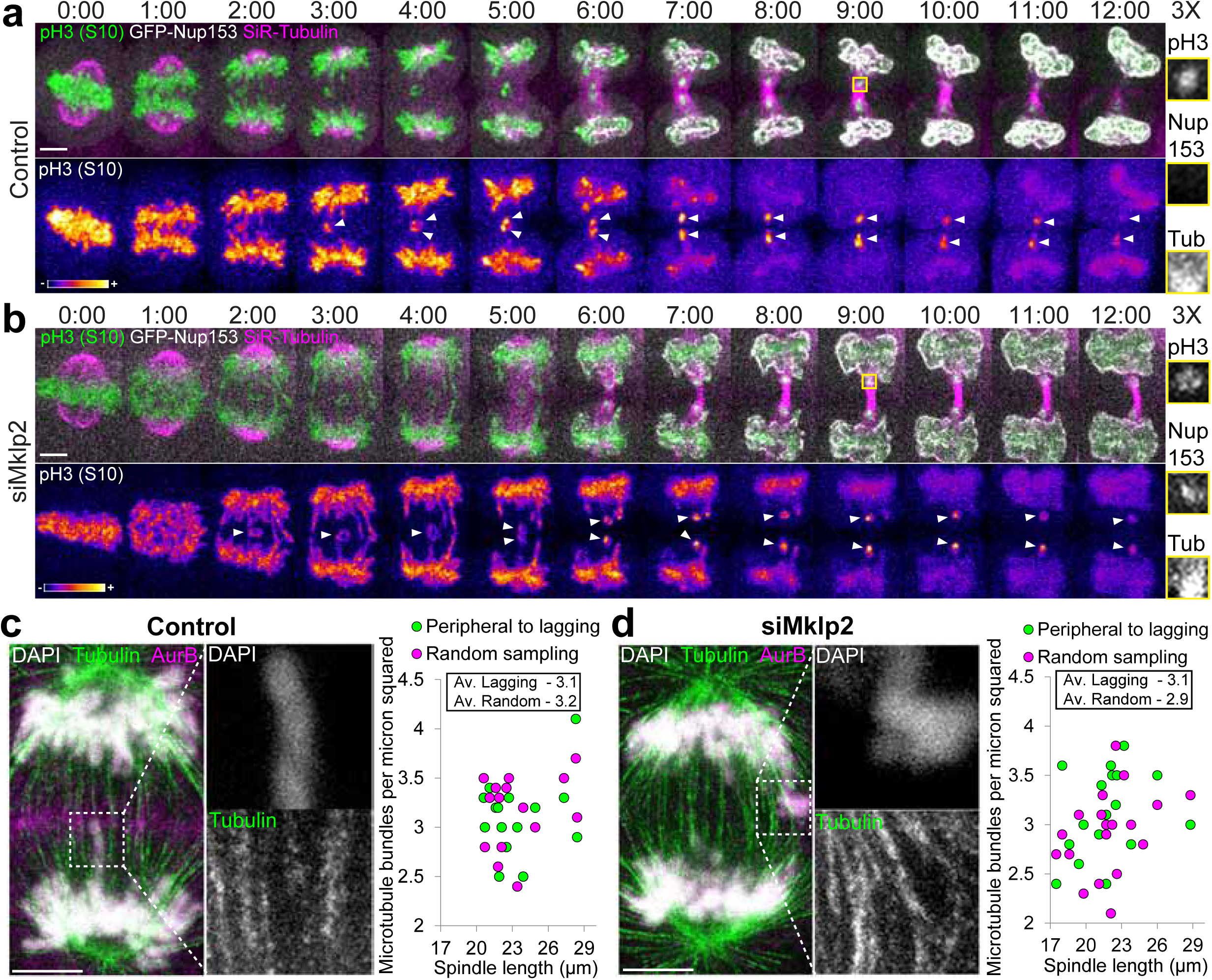
A midzone-based Aurora B activity gradient prevents premature Nup153 recruitment to lagging chromosomes independently of microtubules. Representative examples of **a,** Control and **b,** Mklp2-depleted cells displaying pH3 Ser10 (green), GFP-Nup153 (white) and SiR-tubulin (magenta). Time=min:sec. pH3 Ser10 alone (FIRE LUT) is shown below each cell for better visualization. White arrowheads track lagging chromosomes until they form MN. Insets show 3X magnification of selected regions with lagging chromosomes (grayscale for single channels of pH3, Nup153 and Tubulin). Representative CH-STED images of fixed **c,** control and **d,** Mklp2-depleted anaphase cells displaying DNA (white), α-tubulin (green) and Aurora B (magenta). Insets show 3X magnification of selected regions with lagging chromosomes (grayscale for single channels of DNA and Tubulin). Quantifications on the right show the average number of MT bundles around lagging chromosomes (Peripheral to lagging) and at randomly selected regions on the mitotic spindle (Random sampling). Each dot represents the average measurement for one cell and the numbers shown within the black rectangle represent the averages for all cells. Scale bars = 5 μm.

## Discussion

Taken together, our data demonstrate that a midzone Aurora B phosphorylation gradient mediates an anaphase surveillance mechanism that integrates error correction by mechanical transduction of spindle forces with spatial control of NER on lagging chromosomes, to protect against MN formation in human cells. The immediate implication of our findings is that human cells normally progress through anaphase with one or more lagging chromosomes, but these are actively “checked” and corrected during anaphase to avoid MN formation, as predicted by the ‘chromosome separation checkpoint’ hypothesis (Afonso et al., 2014; Maiato et al., 2015). Indeed, merotelic attachments are still detectable during metaphase (Knowlton et al., 2006) and their correction is promoted by experimentally increasing metaphase duration by two hours (Cimini et al., 2003), suggesting that few mitotic errors normally escape the surveillance of the SAC that is satisfied within ∼20 minutes upon attachment of the last kinetochore to the spindle (Rieder et al., 1995). What our results uncover is that this scenario is just the tip of the iceberg and many more errors are revealed once the dependence that characterizes all checkpoints is relieved (Hartwell and Weinert, 1989). In agreement, we observe both a significant increase of anaphase cells with lagging chromosomes, as well as an increase in the number of lagging chromosomes per anaphase cell when Aurora B activity and midzone localization is specifically inhibited after anaphase onset. Thus, this increase in lagging chromosomes cannot be due to errors that formed *de novo* after anaphase onset, but rather due to compromised error correction during anaphase, unveiling that SAC-invisible errors are more frequent than previously anticipated.

Error correction during anaphase is known to require the mechanical action of spindle forces that ultimately resolve merotelic attachments (Cimini et al., 2004; Cimini et al., 2003). Here we show that, in human cells, these forces rely on a midzone-based Aurora B phosphorylation gradient that assists kinesin-5-mediated spindle elongation during anaphase. Because Aurora B activity at KTs decreases as a function of chromosome separation from the spindle midzone (Fuller et al., 2008), the proximity to the spindle midzone where Aurora B activity is maximal might locally stabilize KT-MT attachments necessary for efficient mechanical transduction of spindle forces involved in the resolution of chromosome segregation errors during anaphase. Consistent with this hypothesis, molecular perturbation of either Aurora B or its substrates within the KMN network disrupted the stabilization of KT-MT attachments on lagging chromosomes and resulted in MN formation.

According to the ‘chromosome separation checkpoint’ hypothesis (Afonso et al., 2014; Maiato et al., 2015), an anaphase error correction mechanism would not be possible without a coordinated delay in the completion of NER on lagging chromosomes to prevent MN formation. A chromatin-localized Aurora B-mediated delay in NER has previously been implicated in the prevention of MN formation from late segregating “acentric” chromosomes in *Drosophila* neuroblasts by excluding HP1 from heterochromatin (Warecki and Sullivan, 2018). In this case, late segregating “acentric” chromosomes remain tethered with the segregated chromosome mass and reintegrate the main nuclei by passing through Aurora B-dependent channels in the NE (Karg et al., 2015). Here we show that in human cells, lagging chromosomes, but not DNA bridges, are actively corrected by a midzone-dependent Aurora B activity gradient that locally prevents the completion of NER. Although the local inhibition of NER on both lagging chromosomes and DNA bridges relies on a midzone phosphorylation gradient, the resolution of bridges might instead rely on Aurora B activity on chromatin, as proposed for tethered “acentrics” in *Drosophila* neuroblasts (Karg et al., 2015). The specific effect of the midzone Aurora B phosphorylation gradient on the resolution of lagging chromosomes might reflect the nature of the underlying correction mechanism by spindle forces and may be facilitated by localized delays in the completion of NER in the midzone-facing side of the main segregating chromosome mass (Gerlich and Ellenberg, 2003; Lu et al., 2011) to allow reintegration of lagging chromosomes into the main nuclei.

The existence of an active checkpoint-based mechanism that actively monitors chromosome position during anaphase has been recently challenged, and midzone MTs were proposed to act as a selective physical barrier that irreversibly prevents the specific recruitment of non-core NE components to anaphase lagging chromosomes, independently of Aurora B activity and its midzone localization (Liu et al., 2018). Completion of NER on lagging chromosomes would therefore be strictly dependent on the disassembly of midzone MTs as cells exit mitosis (Liu and Pellman, 2020), with irreversible NE defects on MN emerging as an unsupervised pathological condition that inevitably links mitotic errors to chromothripsis. This idea is supported by elegant correlative live-cell and electron microscopy experiments suggesting that MTs might block NE assembly (Haraguchi et al., 2008) and the observation that MT stabilization near chromosomes irreversibly inhibited the recruitment of non-core NE components in *Xenopus* egg extracts (Xue et al., 2013). Although the two models are not mutually exclusive, the interdependence between microtubules and Aurora B activity makes it difficult to experimentally separate their respective roles in NE formation. For instance, MT stabilization near chromosomes was shown to be dependent on Aurora B (Xue et al., 2013) and MTs increase Aurora B kinase activity towards several microtubule-associated substrates *in vitro* (Noujaim et al., 2014). Because Aurora B activity on chromosomes and spindle midzone MTs decreases as a function of chromosome separation during anaphase in live human cells (Fuller et al., 2008; Tan and Kapoor, 2011), it is conceivable that MTs prevent the recruitment of non-core NE proteins to lagging chromosomes by regulating the Aurora B-dependent phosphorylation of specific substrates on chromatin or the NE (Afonso et al., 2017). Noteworthy, the proposal that midzone MTs, rather than an Aurora B activity gradient, physically prevent the completion of NER on lagging chromosomes was based on fixed-cell experiments in which RPE1 cells depleted of Kif4A, a chromokinesin that is recruited to overlapping MT bundles at the spindle midzone by PRC1 (Bieling et al., 2010; Kurasawa et al., 2004; Subramanian et al., 2013), but not cells depleted of Mklp2, showed premature accumulation of non-core NE components to lagging chromosomes induced after nocodazole treatment/washout (Liu et al., 2018). These findings are at odds with the live-cell data presented in the present study and are difficult to reconcile with the fact that Mklp2 is required for Kif4A accumulation at the spindle midzone (Nunes Bastos et al., 2013) (see also Fig. S3e-g) and known roles of Kif4A in Aurora B recruitment to this location (Kurasawa et al., 2004). Not less important, our close inspection of spindle midzone MTs in the absence of Mklp2 by super-resolution CH-STED microscopy (Pereira et al., 2019) failed to detect significant differences in MT density in the vicinity of anaphase lagging chromosomes, and similar findings have been reported after expansion microscopy analysis of human cells depleted of PRC1, which prevents Kif4A recruitment to midzone MTs (Vukušić et al., 2019). One possibility is that Kif4A spatially controls the completion of NER on lagging chromosomes by regulating Condensin I-mediated chromosome condensation (Afonso et al., 2014; Mazumdar et al., 2004; Poonperm et al., 2017; Samejima et al., 2012; Takahashi et al., 2016). Furthermore, as we show here, the perturbation of the Aurora B gradient due to unintended effects caused by a prolonged mitotic arrest and/or the presence of residual intracellular nocodazole due to incomplete washout cannot be excluded.

Key to this puzzle is whether one can devise assays that can experimentally separate Aurora B activity from its MT dependency in the control of NER. Complete MT depolymerisation with colchicine in living *Drosophila* cells in culture, followed by acute pharmacological inhibition of Aurora B was sufficient to trigger NER (Afonso et al., 2014). One caveat though, is that acute Aurora B inhibition also abrogates the SAC (Moutinho-Pereira et al., 2013; Santaguida et al., 2011) and thus NER may just have occurred as a consequence of mitotic exit. Nevertheless, recent independent work has shown that isolated MN generated in the absence of MTs lack functional NEs that become prone to rupture, suggesting that MTs do not account for the NE defects commonly observed on MN (Kneissig et al., 2019). On the other hand, taxol treatment, together with Aurora B inhibition, favored the idea that midzone MTs, rather than Aurora B activity, prevent the recruitment of non-core NE proteins to chromatin (Liu et al., 2018). However, Aurora B inhibition in these experiments was performed only 4 minutes after anaphase onset, and we show here that Nup153 and LBR recruitment to lagging chromosomes after Mklp2 depletion, as well as Aurora B transport to the spindle midzone in control cells, takes less than 2 minutes after anaphase onset. Thus, Aurora B inhibition later in anaphase might not be sufficient to revert the phosphorylation gradient that spatially controls NER. Lastly, our high-resolution live-cell assay where the Aurora B phosphorylation gradient on chromosomes was simultaneously monitored with the recruitment of non-core NE components and the distribution of mitotic spindle MTs, unequivocally demonstrates that a midzone Aurora B phosphorylation gradient on chromosomes, rather than midzone MTs, delays the completion of NER on lagging chromosomes in human cells. This conclusion gains weight after the recent discovery that Aurora B-mediated phosphorylation of chromatin-associated cGAS prevents its premature activation during mitosis (Li et al., 2021), raising the exciting possibility that an Aurora B-dependent ‘chromosome separation checkpoint’ links mechanisms of MN prevention, with those involved in the clearance of micronucleated cells. Overall, our findings unveil a new layer of protection against genomic instability operating during late mitosis in human cells, awareness of which will be critical for the rational design of MN-targeting therapies.

### Author contributions

Conceptualization, Supervision, Project Administration and Funding acquisition (HM); Methodology (BO, FDS, AMG; LTF, ACF); Investigation, Formal Analysis and Validation (BO, FDS, AMG); Visualization (BO, FDS, AMG, HM); Writing – Original Draft (HM, BO); Writing – Review and Editing (BO, FDS, AMG, LTF, HM).

## Supporting information

Graphical Abstract

## Acknowledgments

We thank António Pereira and Ana Almeida for assistance with CH-STED, Olga Afonso and Ariana Jacome for the generation of lentiviral vectors, Cristina Ferrás and Marco Novais-Cruz for drawing attention to the effects of nocodazole treatment/washout on the Aurora B phosphorylation gradient, current and former lab members for suggestions and Jonathan Higgins for communicating unpublished results. This work was funded by the European Research Council (ERC) consolidator grant CODECHECK, under the European Union’s Horizon 2020 research and innovation programme (grant agreement No 681443).

## Supplemental Figure Legends

**Fig. S1 (Related to Fig. 1) Inhibition of Aurora B activity, but not midzone localization, during anaphase increases the formation of chromosome bridges. a**, Selected timeframes from RPE1 cells expressing H2B-GFP (inverted, grayscale) treated under the specified conditions. Scale bar = 5 µm. **b**, Frequency of U2OS and RPE1 anaphase cells with DNA bridges under the specified conditions. For U2OS cells (Control n=198, 1 µM ZM447439 (ZM) n=29, 10 µM Paprotrain (Papro) added in Metaphase (Meta) n=33, 10 µM Paprotrain (Papro) added in Anaphase (Ana) n=45, siMklp2 n=43) and for RPE1 cells (Control n=136, 1 µM ZM447439 (ZM) n=19, 2 µM ZM447439 (ZM) n=18, 10 µM Paprotrain (Papro) added in Metaphase (Meta) n=27, 10 µM Paprotrain (Papro) added in Anaphase (Ana) n=21, siMklp2 n=27). **c**, Frequency of micronuclei formation and lagging chromosomes that result in micronuclei in U2OS and RPE1 cells, with and without Mklp2.

**Fig. S2 (Related to Fig. 1) Titration of ZM447439 and Paprotrain identifies the optimal doses for specifically affecting Aurora B function in U2OS cells. a**, Western blot analysis showing a dose-dependent loss of Aurora activity in the presence of ZM447439. Numbers below indicate the relative amount of protein detected (%), normalized to GAPDH levels and relative to controls. To enhance detection of Aurora A, B and C, all cells were pre-treated with 100 nM Taxol for 19 h (according to manufacturer instructions) to enrich the mitotic population before adding titrated doses of ZM447439. **b**, Quantification of mitotic cells treated with different concentrations of Paprotrain (added either during prometaphase or metaphase) that fail to enter anaphase during imaging. **c**, Representative cell treated with 20 µM Paprotrain before nuclear envelope breakdown (NEBD). **d**, Representative metaphase cell treated with 20 µM Paprotrain during late prometaphase. Note the rapid effects on spindle morphology. **e**, Representative metaphase cell treated with 10 µM Paprotrain. For all time-lapse panels time is shown in h:min:sec where 00:00:00 is the time of Paprotrain addition. Scale bar = 5 µm.

**Fig. S3 (Related to Fig. 2) Efficiency of Mklp2 depletion in U2OS cells. a**, Western blot analysis showing efficient Mklp2 depletion 72 h after transfection. GAPDH was used as a loading control. **b**, Immunofluorescence image of control and siMklp2 cells fixed and stained to reveal DAPI (blue), Mklp2 (green) and INCENP (red). Note that Mklp2 depletion results in INCENP delocalization. Horizontal line-scan averages of the relative levels of Mklp2 and INCENP in **c**, control and **d**, Mklp2-depleted anaphase cells (Control n=15, siMklp2 n=15; error bars represent SEM). **e**, U2OS cells fixed and stained to reveal DAPI (blue), Kif4a (green) and Aurora B (red). Horizontal line-scan averages of the relative levels of Kif4a and Aurora B in **f**, control and **g**, Mklp2-depleted anaphase cells (Control n=10, siMklp2 n=10; error bars represent SEM). Note that Kif4a localizes to chromatin and the midzone in control cells but fails to localize at the midzone in the absence of Mklp2. Scale bars = 5 μm.

**Fig. S4 (Related to Fig. 1) Disruption of Aurora B and/or kinesin-5 function affects spindle elongation velocity and chromosome segregation distance in U2OS cells.** Quantifications of **a**, spindle elongation velocity [Control n=21, siMklp2 n=21, FCPT (100 μM) n=7, ZM (1 μM ZM447439) n=19, Paprotrain (10 μM added in Anaphase) n=19, siMklp2+FCPT (100 μM) n=8 ZM(1 μM)+FCPT(100 μM) n=4] and **b**, chromosome segregation distance, measured 7 min after anaphase onset [Control n=15, siMklp2 n=15, FCPT (100 μM) n=15, ZM (1 μM ZM447439) n=15, Paprotrain (10 μM added in Anaphase) n=15, siMklp2+FCPT (100 μM) n=13; ZM(1 μM)+FCPT(100 μM) n=8].

**Fig. S5 (Related to Fig. 3). Prolonged nocodazole treatment and washout prevents full centromere-to-midzone re-localization of Aurora B and disrupts its activity gradient during anaphase in RPE1 cells. a**, Representative images of RPE1 cells used for quantifications shown in **b**, and **c**. Scale bar = 10 μm. Relative Front/Back pH3(S10) ratio of anaphase cells calculated for **b**, RPE1 cells without segregation errors (Control n=15, NocWO n=16) and **c**, RPE1 cells with segregation errors cells (Control n=2, NocWO n=19) under the specified conditions (magenta and green lines indicate mean±SD). Note that chromosome segregation errors in untreated RPE1 cells are rare.

**Fig. S6 (Related to Fig. 5) Aurora B midzone localization prevents premature Nup153 recruitment on DNA bridges during anaphase.** Representative examples of **a**, a control cell with a DNA bridge that is positive for pH3 and negative for Nup153 and **b**, an Mklp2-depleted cell with DNA bridges that are positive for both pH3 and Nup153 (Time is shown above in min:sec). Cells shown in **a**, and **b**, are U2OS cells stably expressing eGFP-Nup153 (inverted; grayscale) and loaded with Fab311-Cy3, which recognizes endogenous pH3 Ser10 (FIRE LUT). Magenta arrows shown in **b**, highlight chromosome errors that prematurely accumulate Nup153 in siMklp2-depleted cells. Scale bar = 5 μm.

**Fig. S7 (Related to Fig. 6) Midzone Aurora B prevents premature LBR recruitment to lagging chromosomes during anaphase.** Representative examples of **a**, a control cell with lagging chromosomes and **b**, an Mklp2-depleted cell with segregation errors (Time is shown in min:sec). Cells shown in panels **a-b** are U2OS cells stably expressing mRFP-H2B (inverted; grayscale) and transiently expressing eGFP-LBR (FIRE LUT). Red arrows highlight chromosome errors that do not accumulate LBR in control cells. Insets represent single planes from z-stacks of selected time frames for better LBR visualization on missegregating chromosomes. Scale bar = 5 μm. **c**, Quantification of the percentage of anaphase cells with LBR on lagging chromosomes (Control n=16, siMklp2 n=40). **d**, Difference in time (min) between detection of LBR on daughter nuclei and lagging chromosomes (Control n=16, siMklp2 n=40). e, Quantification of the time (min) it takes to detect LBR on daughter nuclei after anaphase onset (Control n=17, siMklp2 n=43).

## Legends for Supplemental Movies S1 to S7

**Movie S1. Aurora B activity and midzone localization during anaphase are essential for mitotic fidelity in U2OS cells.** Spinning-disk confocal time-lapse imaging of U2OS cells stably expressing H2B-GFP (green) and mCherry-α-tubulin (magenta) under the indicated experimental conditions. Cells were imaged every 30 seconds (sec). For ZM447439, Paprotrain and FCPT treatments, 1 μM, 10 μM and 100 μM were added respectively within 60 seconds of anaphase onset. Time is displayed in seconds (sec) and time 0 = Anaphase Onset. Scale bar = 5 μm.

**Movie S2. Aurora B activity and midzone localization during anaphase are essential for mitotic fidelity in RPE1 cells.** Spinning-disk confocal time-lapse imaging of RPE1 cells stably expressing H2B-GFP (inverted grayscale) under the indicated experimental conditions. Cells were imaged every 30 seconds (sec). For ZM447439 and Paprotrain treatments, 2 μM and 10 μM were added respectively within 60 seconds of anaphase onset. Time is displayed in seconds (sec) and time 0 = Anaphase Onset. Scale bar = 5 μm.

**Movie S3. Aurora B fails to localize to the spindle midzone during anaphase in the absence of Mklp2.** Spinning-disk confocal time-lapse imaging of HeLa cells stably expressing LAP-Aurora B (inverted, grayscale) under the indicated experimental conditions. Cells were imaged every 30 seconds (sec). For ZM447429 and Paprotrain treatments, 1 μM and 10 μM were added respectively within 60 seconds of anaphase onset. Time is displayed in seconds (sec) and time 0 = Anaphase Onset. Scale bar = 5 μm.

Movie S4. Aurora B midzone localization during anaphase is required to sustain a spatially regulated phosphorylation gradient during anaphase. Spinning-disk confocal time-lapse imaging of U2OS cells loaded with Fab311-Cy3, that recognizes endogenous pH3 Ser10 (Left panel; FIRE LUT) and stably expressing GFP-Nup153 (right panel; inverted, grayscale) in control and siMklp2-treated cells. Cells were imaged every 30 seconds (sec). Time is displayed in seconds (sec) and time 0 = Anaphase Onset. Scale bar = 5 μm.

**Movie S5. Aurora B midzone localization during anaphase is required to prevent premature Nup153 accumulation on lagging chromosomes.** Spinning-disk confocal time-lapse imaging of U2OS cells stably expressing RFP-H2B (left panel; inverted, grayscale) and transiently expressing eGFP-Nup153 (Right panel; FIRE LUT) in control and siMklp2-treated cells. Cells were imaged every 30 seconds (sec). Time is displayed in seconds (sec) and time 0 = Anaphase Onset. Scale bar = 5 μm.

**Movie S6. Aurora B midzone localization during anaphase is required to prevent premature LBR accumulation on lagging chromosomes.** Spinning-disk confocal time-lapse imaging of U2OS cells stably expressing RFP-H2B (left panel; inverted, grayscale) and transiently expressing eGFP-LBR (Right panel; FIRE LUT) in control and siMklp2-treated cells. Cells were imaged every 30 seconds (sec). Time is displayed in seconds (sec) and time 0 = Anaphase Onset. Scale bar = 5 μm.

**Movie S7 – Aurora B midzone localization is required for spatial control of NER independently of microtubules.** Left panels represent spinning-disk confocal time-lapse imaging of U2OS cells stably expressing eGFP-Nup153 (white), loaded with Fab311-Cy3, that recognizes endogenous pH3 Ser10 (green) and stained with SiR-Tubulin (magenta) in control and siMklp2-treated cells. Middle panels show single channel images of pH3 Ser10 (FIRE LUT) and panels on the right show single channel images of GFP-Nup153 (inverted, grayscale). Cells were imaged every 30 seconds (sec). Time is displayed in seconds (sec) and time 0 = Anaphase Onset. Scale bar = 5 μm.

## Online content

Any methods, additional references, supplemental data and movies, acknowledgements, details of author contributions and competing interests; and statements of data availability.

## STAR Methods

### Cell Culture

All cell lines were cultured at 37°C in 5% CO_2_ atmosphere in Dulbecco’s modified medium (DMEM; Gibco™, Thermofisher) containing 10% fetal bovine serum (FBS; Gibco™, Thermofisher). U2OS parental and H2B-eGFP/mCherry-α-tubulin U2OS cells were kindly provided by S. Geley (Innsbruck Medical University, Innsbruck, Austria). hTERT-RPE1 (RPE1) parental (ATCC® CRL-4000™) and HeLa LAP-Aurora B cells kindly provided by Ben Black (U. Pennsylvania, PA, USA). Stable mRFP-H2B U2OS and H2B-eGFP/mCherry-α-tubulin RPE1 cells were generated by lentiviral transduction as previously described (Ferreira et al., 2018). Stable eGFP-Nup153 U2OS cells were generated through transfection of an eGFP-Nup153 plasmid using Lipofectamine™ 2000 according to the manufacturer instructions. Single cell clones were expanded and assayed for correct localization and moderate expression levels. One clone was processed using Fluorescence Activated Cell Sorting (FACS; FACS Aria II) and selected for use as a stable cell line throughout this study.

### Constructs and Transfections

To express fluorescently-tagged proteins, cells were transfected using Lipofectamine™ 2000 (Thermofisher) with 1 μg of each construct. Plasmids used were: pEGFP(C3)-Nup153 (Addgene #64268); eGFP-LBR and eGFP-Nup35, kindly provided by Tokuko Haraguchi (Advanced ICT Research Institute Kobe, NICT, Kobe, Japan) (Haraguchi et al., 2008). To deplete Mklp2 by RNA interference (RNAi), cells were plated at 40-50% confluence onto 22 x 22 mm No. 1.5 glass coverslips and cultured for 12 h in DMEM supplemented with 10% of FBS before transfection. Cells were starved in serum-free media (Opti-MEM™, Thermofisher) by incubating 1 h before RNAi transfection. RNAi transfection was performed using Lipofectamine™ RNAiMAX reagent (Thermofisher) with 25 nM of validated siRNA oligonucleotides against human Mklp2: 5’-AACGAACUGCUUUAUGACCUA-3’ (Sigma Aldrich) and control (scramble) siRNA 5’-UGGUUUACAUGUCGACUAA-3’ (Sigma Aldrich), diluted in serum-free media (Opti-MEM™, Thermofisher). Untreated, mock transfected or scramble siRNA results were indistinguishable and were therefore referred to as Control. Depletion of Mklp2 was maximal at 72h after siRNA transfection and all of the analysis was performed at 72h.

### Western Blotting

Cell extracts were collected after trypsinization and centrifuged at 1200 rpm for 5 min, washed and re-suspended in Lysis Buffer (NP-40, 20 nM HEPES/KOH pH 7.9 ; 1 mM EDTA pH 8; 1 mM EGTA; 150 mM NaCl; 0.5% NP40; 10% glycerol, 1:50 protease inhibitor; 1:100 Phenylmethylsulfonyl fluoride). The samples were then flash frozen in liquid nitrogen and kept on ice for 30 min. After centrifugation at 14000 rpm for 10 min at 4°C the supernatant was collected and protein concentration determined by the Bradford protein assay (Bio-Rad). The proteins were run on SDS-PAGE gels (50 μg per lane) and transferred at constant amperage for 10 min using the iBlot Gel Transfer Device (Thermo Scientific), to a nitrocellulose Hybond-C membrane. Membranes were then blocked with 5% Milk in TBS with 0.1% Tween-20 (TBS-T) for 1 h at room temperature. The primary antibodies used were: anti-rabbit Mklp2/Kif20A (Bethyl Laboratories; 1:5000); anti-rabbit phospho Aurora A/B/C (T288/232/398; Cell Signaling; 1:2000); anti-rabbit Phospho-Histone H3 Ser10 (Monoclonal; D2C8; Cell Signaling Technology; 1:5000); anti-mouse GAPDH (Monoclonal; Proteintech; 1:40000). All primary antibodies were incubated overnight at 4°C with shaking. After three washes in TBS-T the membranes were incubated with the secondary antibody for 1-3 h at room temperature. The secondary antibodies used were anti-mouse-HRP (Jackson ImmunoResearch) and anti-rabbit-HRP (Jackson ImmunoResearch) at 1:5000. After several washes with TBS-T, the detection was performed with Clarity Western ECL Substrate (Bio-Rad).

### Time-lapse spinning-disk confocal microscopy

For time-lapse microscopy, cells were plated onto 22 x 22 mm (No. 1.5) glass coverslips (Corning) and cell culture medium was changed to DMEM with 10% FBS (without phenol red and supplemented with HEPES buffer) 6-12 h before mounting. Coverslips were mounted onto 1-well Chamlide CMS imaging chambers (Microsystem AB; Sweden) immediately before imaging. All live-cell imaging experiments were performed at 37 °C using temperature-controlled Nikon TE2000 microscopes equipped with a modified Yokogawa CSU-X1 spinning-disc head (Yokogawa Electric), an electron multiplying iXon+ DU-897 EM-CCD camera (Andor) and a filter-wheel. Three laser lines were used for excitation at 488, 561 and 647nm. All live cell imaging experiments were performed using an oil-immersion 100x 1.4 NA Plan-Apo DIC (Nikon), with the exception of the experiments conducted with co-expression of Fab311-Cy3 and eGFP-Nup153, that were performed using an oil-immersion 60x 1.4 NA Plan-Apo DIC (Nikon). All image acquisition was controlled by NIS Elements AR software. Images were collected every 30 seconds: 9 x 2 μm z-stacks spanning a total volume of 16 μm

### Immunofluoresence and fixed cell analysis

For immunofluorescence processing, cells were fixed with 4% Paraformaldehyde (Electron Microscopy Sciences) for 10 min followed by extraction with 2 x 5 min washes using 0.5% Triton X-100 (Sigma-Aldrich). Primary antibodies used were: anti-mouse Aurora B (Monoclonal AIM-1; BD Biosciences; 1:250); anti-mouse INCENP (Monoclonal; Santa Cruz Biotechnology; 1:50); anti-rabbit Mklp2/Kif20A (Bethyl Laboratories; 1:2000); anti-rabbit Phospho-Histone H3 Ser10 (Monoclonal; D2C8; Cell Signaling Technology; 1:5000), anti-rabbit Kif4a (Thermofisher; 1:1000). Secondary antibodies used were Alexa Fluor 488 (Themofisher; 1:2000) and Alexa Fluor 568 (Themofisher; 1:2000). DNA was counterstained with 1 μg/mL DAPI (4’,6’-diamino-2-fenil-indol; Sigma-Aldrich) and mounted onto glass slides with 20 mM Tris pH8, 0.5 N-propyl gallate and 90% glycerol. 3D wide-field images were acquired using an AxioImager Z1 (100x Plan-Apochromatic oil differential interference contrast objective lens, 1.46 NA, ZEISS) equipped with a CCD camera (ORCA-R2, Hamamatsu) operated by Zen software (ZEISS). Blind deconvolution of 3D image data sets was performed using Autoquant X software (Media Cybernetics).

### High-content live-cell microscopy screening

Hela cells stably expressing GFP-H2B/mRFP-α-tubulin were plated onto 96-well plate in DMEM supplemented with 5% FBS and after 1 h transfected with siRNA

oligonucleotides against human Nuf2:AAGCAUGCCGUGAAACGUAUA[dT][dT],

Ndc80:GAAUUGCAGCAGACUAUUA[dT][dT],

Spc24:CUCAACUUUACCACCAAGUUA[dT][dT],

Spc25:CUGCAAAUAUCCAGGAUCU[dT][dT],

KNL1:GCAUGUAUCUCUUAAGGAA[dT][dT],

Mis12:GAAUCAUAAGGACUGUUCA[dT][dT],

Dsn1:GUCUAUCAGUGUCGAUUUA[dT][dT],

Nsl1:CAUGAGCUCUUUCUGUUUA[dT][dT],

Aurora B: AACGCGGCACUUCACAAUUGA[dT][dT], at a final concentration of 50 nM. Transfections were performed using Lipofectamine RNAiMAX in Opti-MEM medium (both from Thermo Fisher Scientific) according to the manufacturer’s instructions. Transfection medium was replaced with complete medium after 6 h. Phenotypes were analyzed and quantified between 24 to 72 h later depending on RNAi depletion efficiency as monitored by the percentage of mitotic cells and by western botting (data not shown). For time-lapse microscopy acquisition, cell culture medium was changed to DMEM with 10% FBS without phenol red and supplemented with HEPES buffer) 6-12 h before acquisition. The image acquisition was performed in an INCell Analyzer 2000 (GE Healthcare, Chicago, IL, USA) with a Nikon 20x/0.45 NA Plan Fluor objective according to manufacturer instructions. Images were collected every 10 min for 72 h.

### Stimulated Emission Depletion (STED) microscopy and sample preparation

For the best visualization of microtubules, cells were fixed with a solution of 4% Paraformaldehyde (Electron Microscopy Sciences) with 0.25% Glutaraldehyde (Electron Microscopy Sciences) for 10 min, followed by quenching of autofluorescence using 0.1% Sodium Borohydride (Sigma-Aldrich) for 10 min. Cells were then permeabilized by washing 2 x 5 min with PBS 0.5% Triton X-100 (Sigma-Aldrich). Primary antibodies used were: mouse anti-Aurora B (Monoclonal AIM-1; BD Biosciences; 1:250); rat monoclonal anti-tyrosinated α-tubulin clone YL1/2 (Bio-Rad, 1:100). STAR-580 (1:100, Abberior), and STAR-RED (1:100, Abberior) were used as secondary antibodies, and DNA was counterstained with 1 µg/ml DAPI (Sigma-Aldrich). For Coherent-Hybrid STED (CH-STED) imaging (Pereira et al., 2019), an Abberior ’Expert Line’ gated-STED microscope was used, equipped with a Nikon Lambda Plan-Apo 1.4 NA 60x objective lens. The depletion beam was generated by a bivortex phase mask (radii ratio = 0.88) to improve suppression of background fluorescence in comparison to conventional 2D-STED. This performance stems from the capacity of a bivortex to generate a depletion dip in 3D that, unlike z-STED or 2D+z-STED combinations, preserves resilience to spherical aberration. All acquisition channels (confocal and STED) were performed using a 0.8 Airy unit pinhole. A time-gate threshold of 500 ps was applied to the STED channel.

### Drug treatments

For live visualization of the mitotic spindle in eGFP-Nup153 U2OS cells, SiR-Tubulin (50 nM; Spirochrome) (Lukinavicius et al., 2014) was added to the culture medium 30 min-1 h before acquisition. For acute inhibition of Aurora B activity, ZM447439 (Sigma-Aldrich) was used at 1 μM U2OS and 2 μM for RPE1 cells. For acute inhibition of Aurora B midzone relocalization, Paprotrain (Tocris Bioscience) was used at 10 μM. For inhibition of spindle elongation, FCPT (kind gift from Tim Mitchison, Harvard University, MA, USA) was used at 100 µM. For all live cell experiments, ZM447439, Paprotrain and FCPT were added within the first min after anaphase onset and DMSO was used as a control. For nocodazole washout experiments to induce lagging chromosomes RPE1 cells were treated with 100 ng/mL Nocodazole (Sigma-Aldrich) for 6 h as previously described (Liu et al., 2018). Nocodazole was washed out by rinsing twice with PBS followed by incubation in fresh DMEM with 10% FBS. Cells were fixed and processed for immunofluorescence 40 min after the Nocodazole was washed out.

### Determination of the frequency of lagging chromosomes forming a micronuclei

The frequency of lagging chromosomes forming micronuclei was determined considering only cells that had 1 or 2 lagging chromosomes in anaphase. Cells with >2 lagging chromosomes were excluded to avoid overestimation. For cells with 1 or 2 lagging chromosomes in anaphase, the percentage of lagging chromosomes that resulted in micronuclei formation for each condition was calculated.

### Quantification of mitotic errors

Mitotic errors were tracked and quantified manually through close inspection of H2B localization in single plane confocal images through the full z-stack. Mitotic errors were divided into 2 main classes: lagging chromosomes or DNA bridges and these were discriminated according to location and morphology associated with H2B localization. Lagging chromosomes retained normal DNA condensation and emerged at different stages during anaphase. Most lagging chromosomes were of transient nature and were only scored if visibly delayed in relation to the main chromosome mass, for at least 2 min (i.e. 4 frames) during anaphase. DNA bridges were characterized by stretches of DNA that connected both daughter nuclei and often displayed aberrant DNA condensation as judged by H2B localization. For lagging chromosomes, DNA bridges and micronuclei, cells were classified as having 1, 2 or >2 events. To determine micronuclei origin, fully formed micronuclei were backtracked to reveal whether these originated from unaligned chromosomes, lagging chromosomes or DNA bridges. Only micronuclei derived from lagging chromosomes were considered in this study.

### Quantification of the Aurora B activity gradient (Front/Back and Lagging/Daughter ratio)

Quantification of the Aurora B gradient in anaphase was performed in both fixed and live cells using phosphorylation of histone H3 at Serine 10 (pH3) as a read-out for Aurora B activity on chromosomes. For Front/Back pH3 gradient measurements, each daughter nuclei was divided into near-equal parts: one closer to the midzone (Front) and the other closer to the pole (Back). Background subtracted mean pixel intensities of these regions were measured using ImageJ software and ratios calculated by dividing the ‘Front’ Mean Pixel Intensity / ‘Back’ Mean Pixel Intensity. For Lagging/Daughter pH3 gradient measurements, background subtracted mean pixel intensities for the lagging chromosome (Lagging) and a corresponding region on daughter nuclei (Daughter) were measured using ImageJ software and ratios calculated by dividing the ‘Lagging’ Mean Pixel Intensity / ‘Daughter’ Mean Pixel Intensity. Note that a ratio close to 1 indicates that there is no difference in pH3 levels throughout the DNA. Conversely, a ratio below 1 indicates that the levels of pH3 are lower on the DNA closer to the midzone.

### Visualization of histone H3 phosphorylation in living cells

For visualization of phosphorylation of histone H3 on Serine 10 in living cells, U2OS cells were loaded with a fluorescently labeled specific Fab antibody fragment that recognizes phosphorylation of Serine 10 on Histone H3 (Fab311-Cy3). U2OS cells were at 80-100% confluence at the time of delivery. For each 22 x 22 mm coverslip, 2 μl of 0.5 mg/ml Fab311-Cy3 were loaded using glass beads (Hayashi-Takanaka et al., 2011) and re-grown in DMEM supplemented with 10% FBS (without phenol red and supplemented with HEPES buffer) for 6-8 h prior to imaging.

### Quantification of spindle elongation, chromosome separation velocity and chromosome <u>separation distance</u>

Spindle elongation velocities were measured from images displaying mCherry-α-tubulin. Centrosome to centrosome distance was measured at anaphase onset and at 3 min after anaphase onset and calculations represent the elongation rate (µm/min) for each spindle. Chromosome separation velocities were measured from images displaying H2B-GFP by calculating chromosome separation distance between 0-3 min after anaphase onset (corresponding to the timeframe with the fastest linear elongation velocity). Chromosome separation distance was calculated by measuring the average distance between each chromosome mass (H2B) 7 min after anaphase onset. Measurements were performed using ImageJ software.

### Statistical Analysis

Quantifications of mitotic errors (i.e. lagging chromosomes, DNA bridges and micronuclei) were analyzed using the Fisher’s exact two-tailed test. Quantifications of chromosome separation velocity, spindle elongation, chromosome segregation distance and dynamics of Nup153 and LBR localization were analyzed using the Mann-Whitney test. Quantifications of dynamics of Aurora B localization were analyzed using a two-tailed t-test. For each graph, where applicable, n.d.= not determined, n.s.= non-significant, *p≤0.05, **p≤0.01 ***p≤0.001 and ****p≤0.0001, unless stated otherwise. Pearson Correlation Coefficient (PCC) was calculated using Excel CORREL function. All results presented in this manuscript were obtained from pooling data from at least 3 independent experiments with the exception of the results presented for depletions of KMN proteins in HeLa cells (siKNL1, siMis12 and siNsl1 = 2 independent experiments; siNuf2, siNdc80, siSpc24 and siSpc25 = 1 experiment).

### Quantification of pH3 and Nup153 in live cells

Quantifications were performed using sum projections of in-focus frames for each channel. For Figs. 4e, and 4f, the values for Nup153 and pH3 were normalized to the maximum (during the first 14 min of anaphase) and to the levels in metaphase, respectively.

## Competing interests

Authors declare no competing interests.

**Resource availability**

## Lead contact

Further information and requests for resources and reagents should be directed to and will be fulfilled by the Lead Contact, Helder Maiato.

## Materials availability

The authors hereby declare that all reagents generated in this study (see Key Resources Table) will be made available upon request.

## Data and code availability

This study did not generate/analyze datasets or code.

