## Supplementary material for "An anaphase surveillance mechanism prevents micronuclei formation from mitotic errors": Figure S1

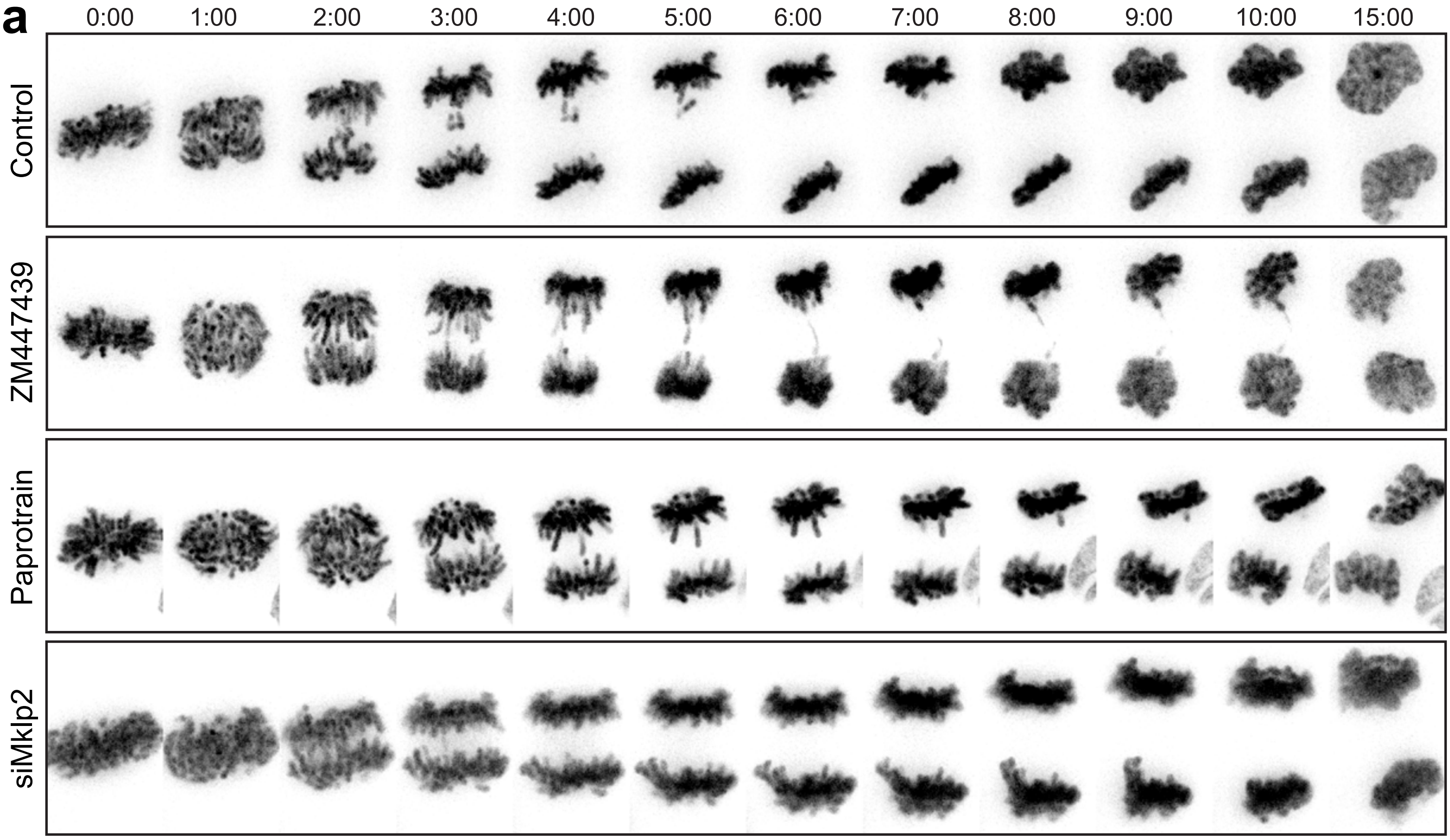


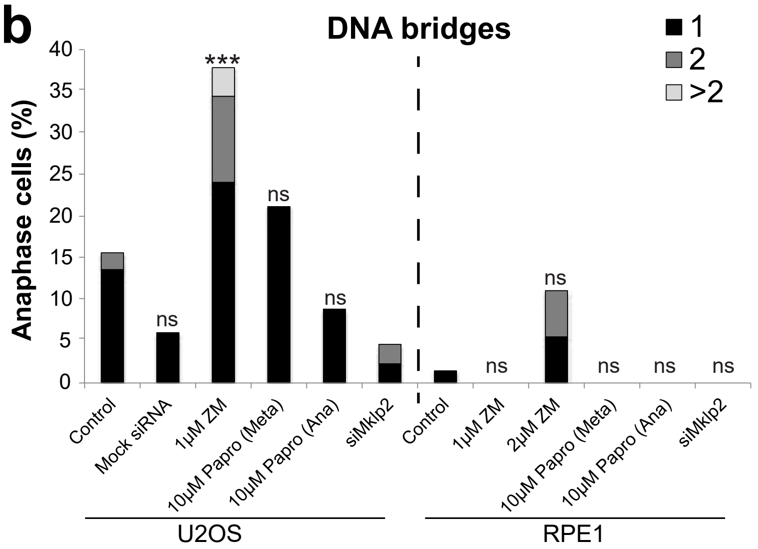


**c**

| | Dividing cells with micronuclei (%) | | Lagging chromosomes that form micronuclei (%) | |
|---|---|---|---|---|
| | **U2OS** | **RPE1** | **U2OS** | **RPE1** |
| Control | 11 | 2 | 14 | 6 |
| siMklp2 | 28 | 11 | 20 | 25 |
