## Supplementary material for "An anaphase surveillance mechanism prevents micronuclei formation from mitotic errors": Figure S2

**a**

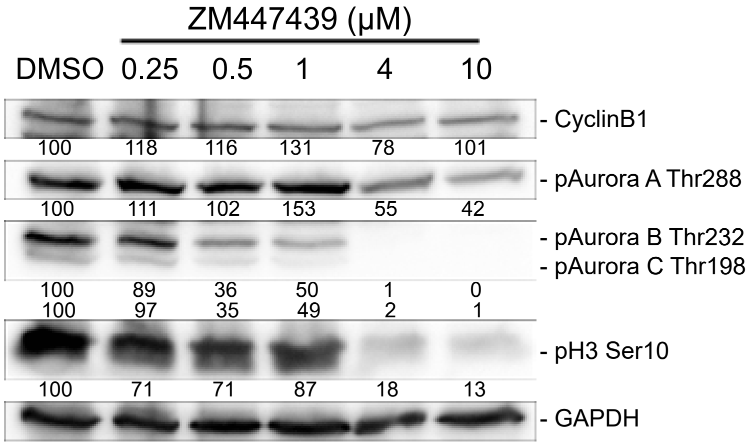


**b**

| Paprotrain | Added during Prometaphase | Added during Metaphase |
|---|---|---|
| | **Cells that do not enter anaphase** | |
| 10 μM | 6/6 (100%) | 21/56 (38%) |
| 20 μM | 4/4 (100%) | n.d. |

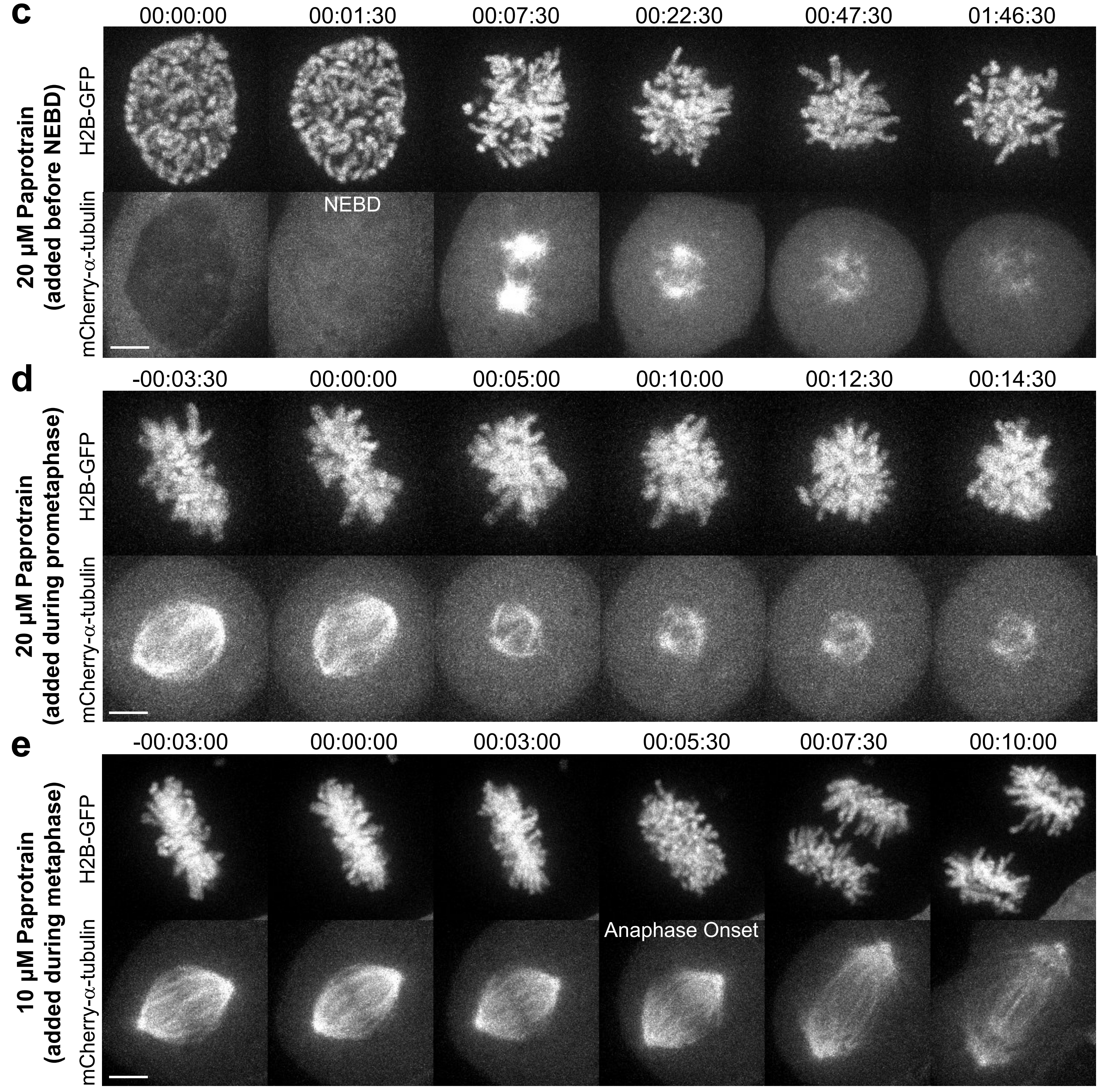
