## Supplementary figures and images for "An anaphase surveillance mechanism prevents micronuclei formation from mitotic errors"

### Figure S3

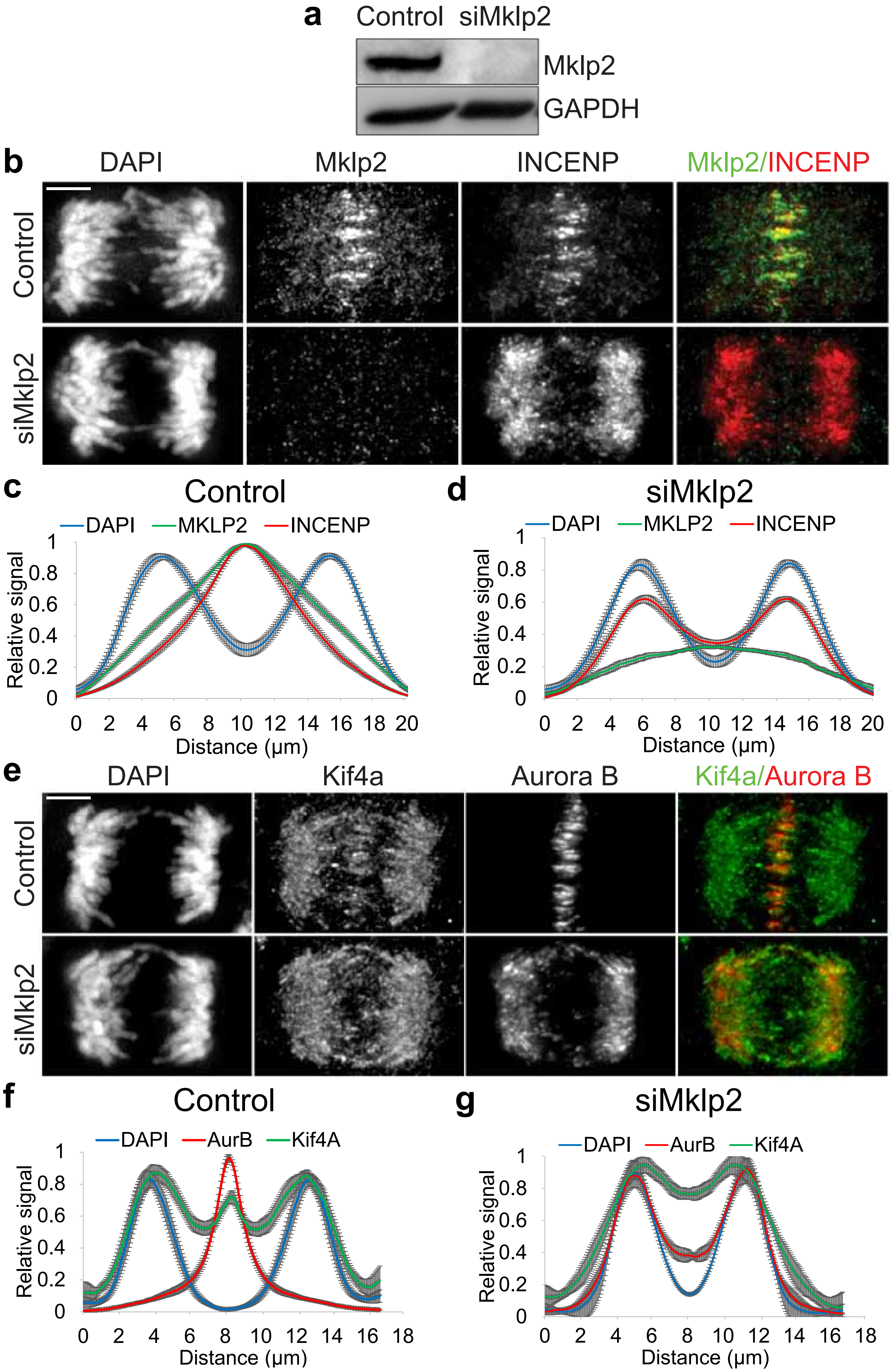

### Figure S4

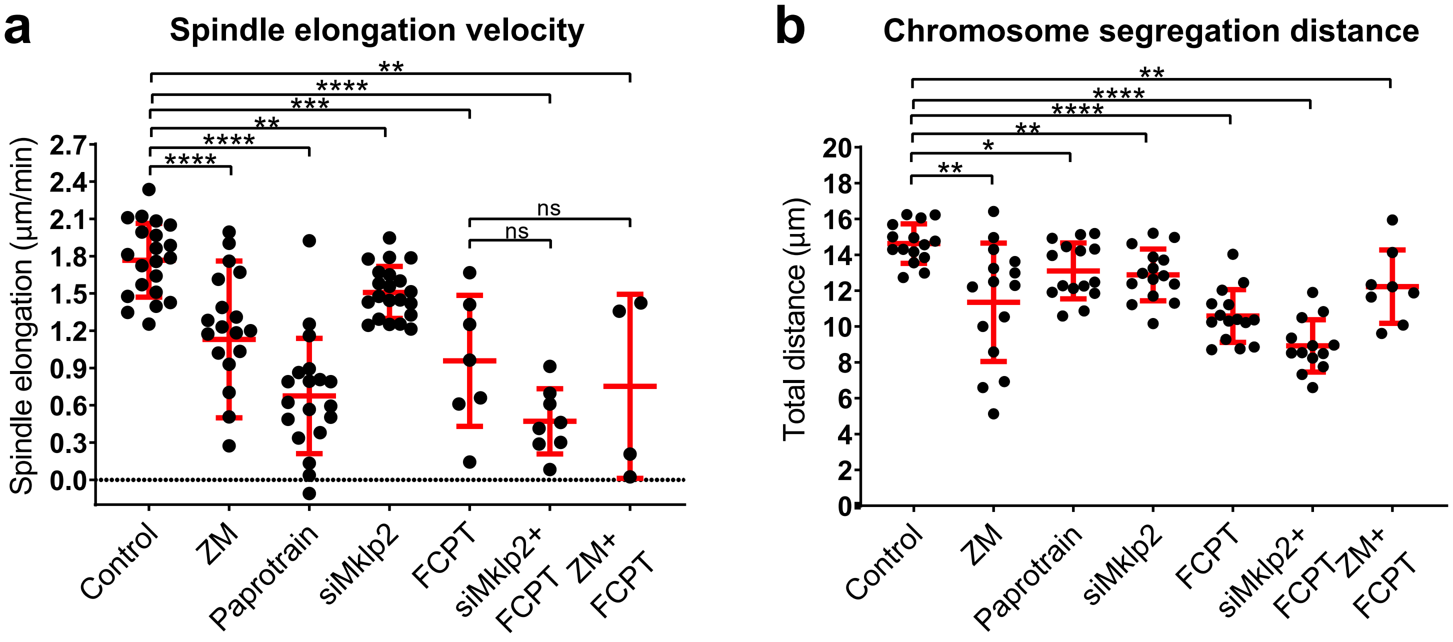

### Figure S5

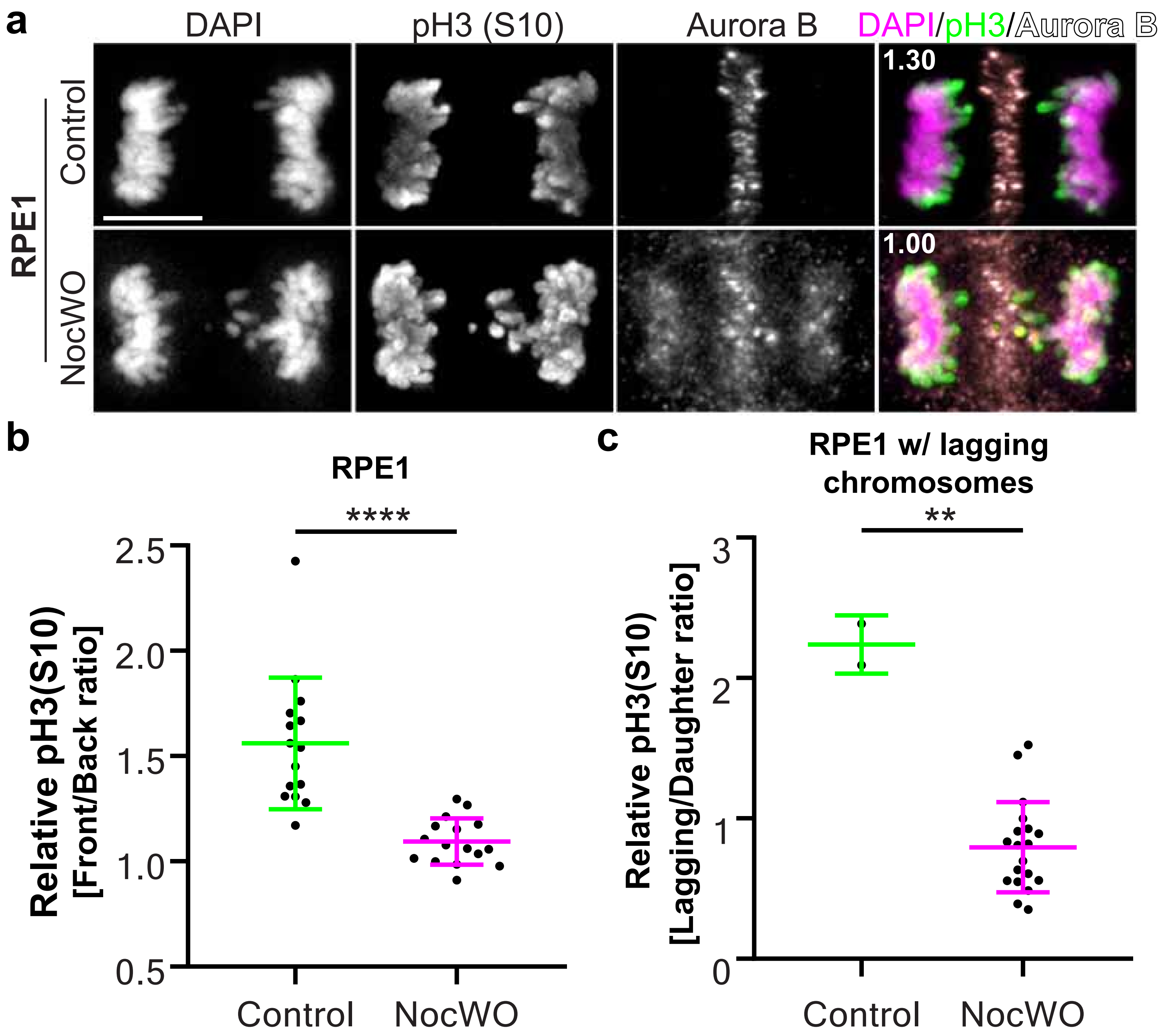

### Figure S6

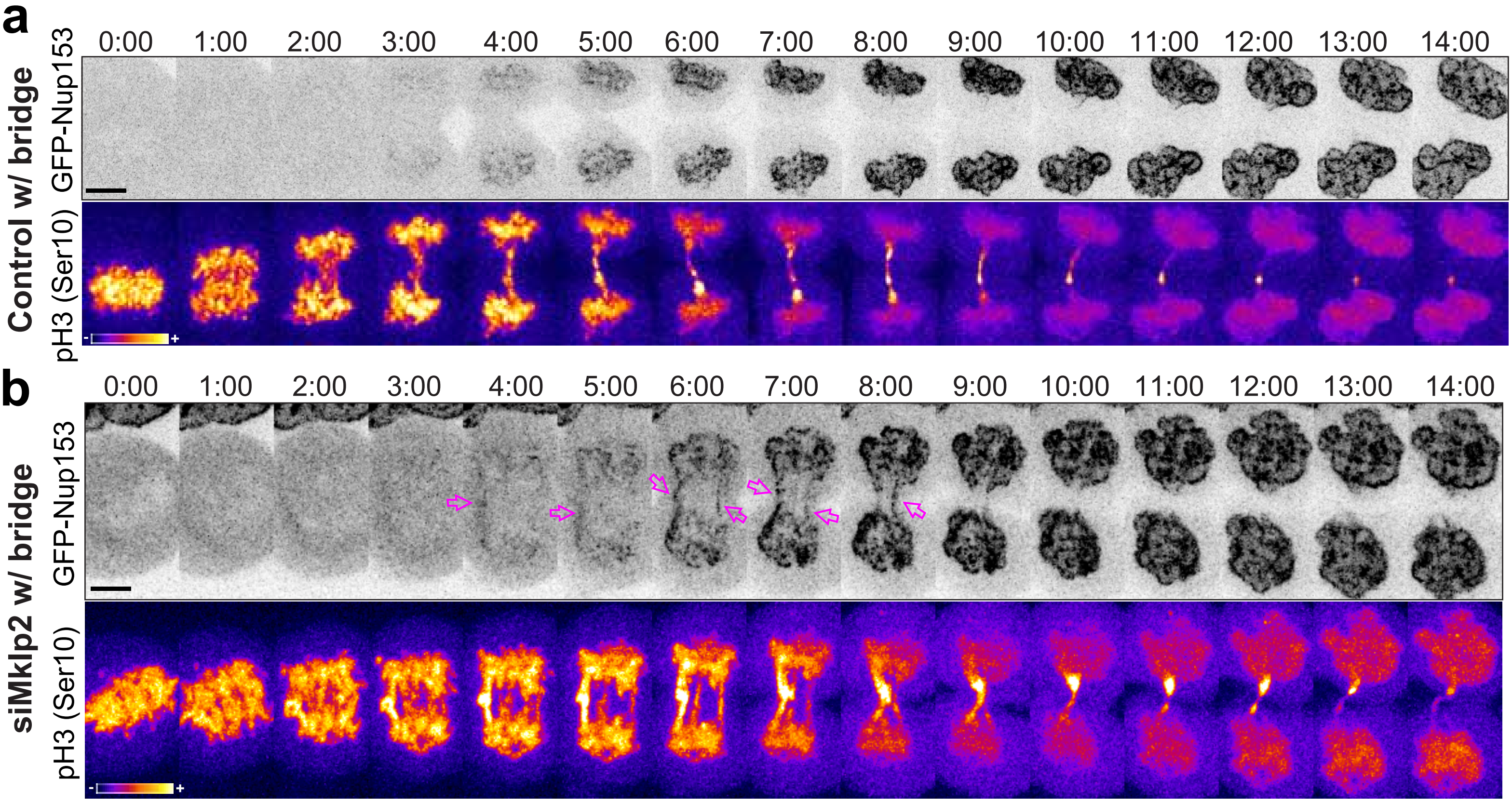

### Figure S7

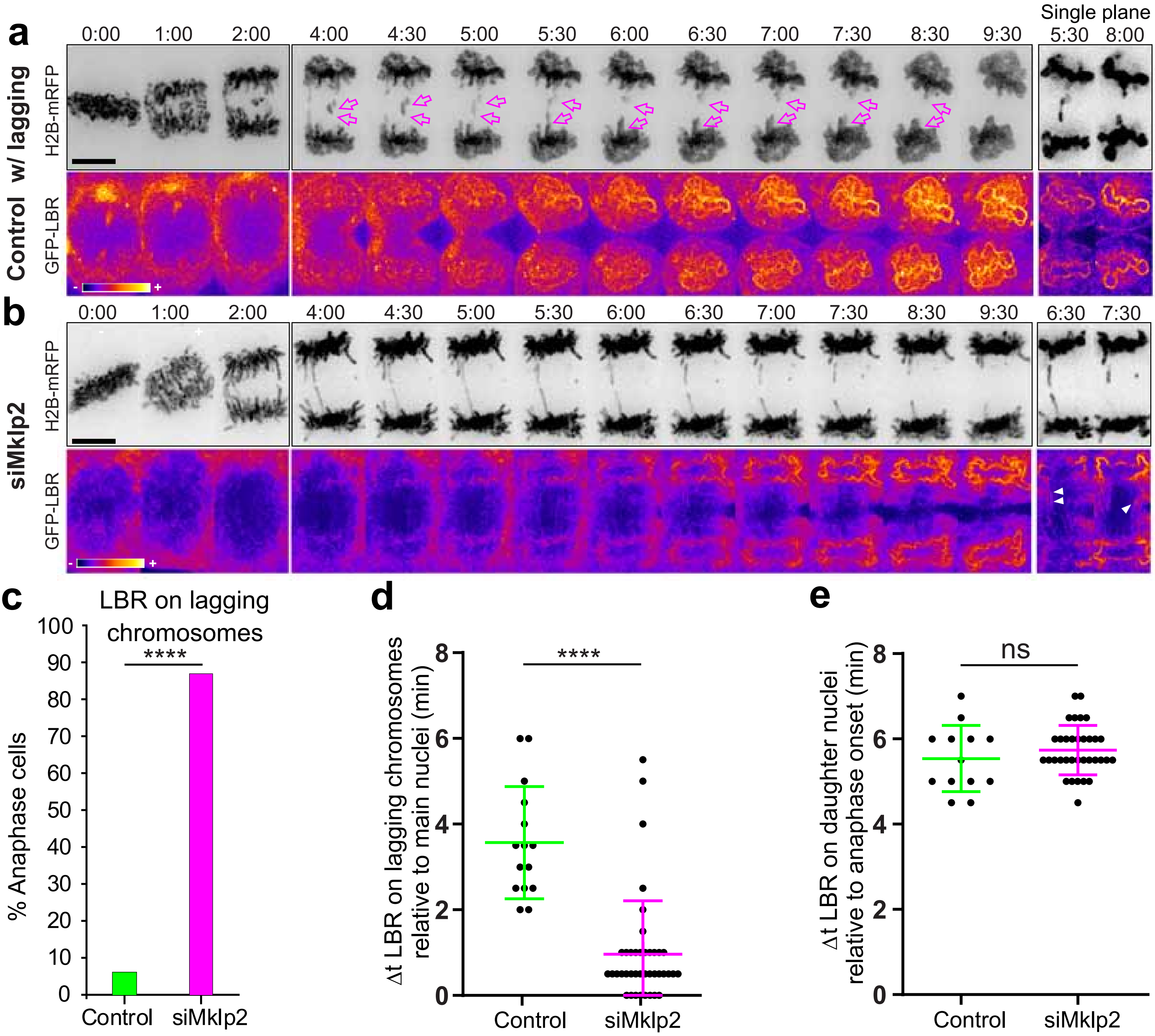

### Graphical Abstract

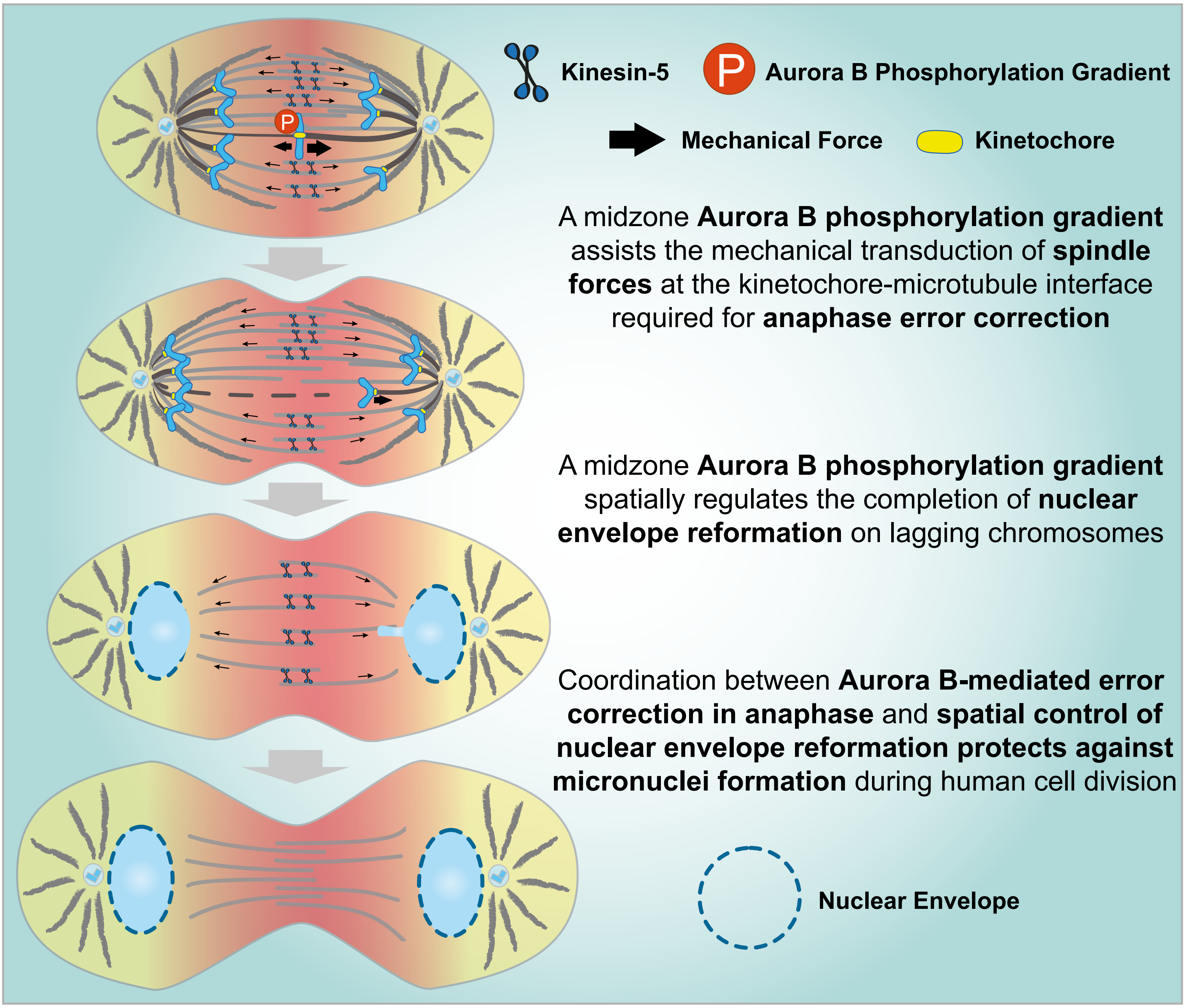
